# Cellular-resolution gene expression mapping reveals organization in the head ganglia of the gastropod, *Berghia stephanieae*

**DOI:** 10.1101/2023.06.22.546160

**Authors:** M. Desmond Ramirez, Thi N. Bui, Paul S. Katz

**Affiliations:** Institute of Neuroscience, University of Oregon, United States; Department of Biology, University of Massachusetts-Amherst, United States; Neuroscience and Behavior Graduate Program, University of Massachusetts-Amherst, United States

## Abstract

Gastropod molluscs such as *Aplysia*, *Lymnaea*, and *Tritonia* have been important for determining fundamental rules of motor control, learning, and memory because of their large, individually identifiable neurons. Yet for the vast majority of gastropod neurons, as well as glia, there are no established molecular markers, limiting the ability to establish brain-wide structure-function relations. Here we combine high-throughput, single-cell RNA sequencing (scRNAseq) with *in-situ* hybridization chain reaction (HCR) in the nudibranch *Berghia stephanieae* to identify and visualize the expression of markers for cell types. Broad neuronal classes were characterized by genes associated with neurotransmitters, like acetylcholine, glutamate, serotonin, and GABA, as well as neuropeptides. These classes were subdivided by other genes including transcriptional regulators and unannotated genes. Marker genes expressed by neurons and glia formed discrete, previously unrecognized regions within and between ganglia. This study provides the foundation for understanding the fundamental cellular organization of gastropod nervous systems.

## Introduction

The central nervous system (CNS), including the brain, contains many different cell types, forming the basis of its complex structure and connectivity, and enabling its sophisticated functions. Cell type diversity in the CNS is reflected in differential gene expression across neurons (***Lein et al., 2007***; ***Vergara et al., 2017***). Analysis of high-throughput single cell RNA sequencing (scRNAseq) allows brain-wide classification of neurons based on gene expression (***Brunet Avalos et al., 2019***; ***Gavriouchkina et al., 2022***; ***Tasic, 2018***; ***Tosches et al., 2018***; ***Styfhals et al., 2022***). The central ring ganglia (CRG) of gastropod molluscs, such as *Aplysia*, *Lymnaea*, and *Tritonia*, contain large neurons, some of which can be individually identified based on immunohistochemistry (IHC), neuroanatomy, and neurophysiology (***Katz and Quinlan, 2019***; ***Leonard, 2000***). These same characteristics can even be used to identify homologous neurons across species (***Croll, 1987***; ***Newcomb et al., 2012***). However, these approaches are generally limited to the small number of large neurons that are impaleable with intracellular microelectrodes and by the availability of antibodies; they do not easily scale up to the thousands of small neurons in the CRG or the even smaller neurons in peripheral ganglia. We combined scRNAseq with *in-situ* hybridization chain reaction (HCR) to obtain a more complete atlas of neurons in head ganglia of the nudibranch, *Berghia stephanieae* (Fig. 1).

**Figure 1.**
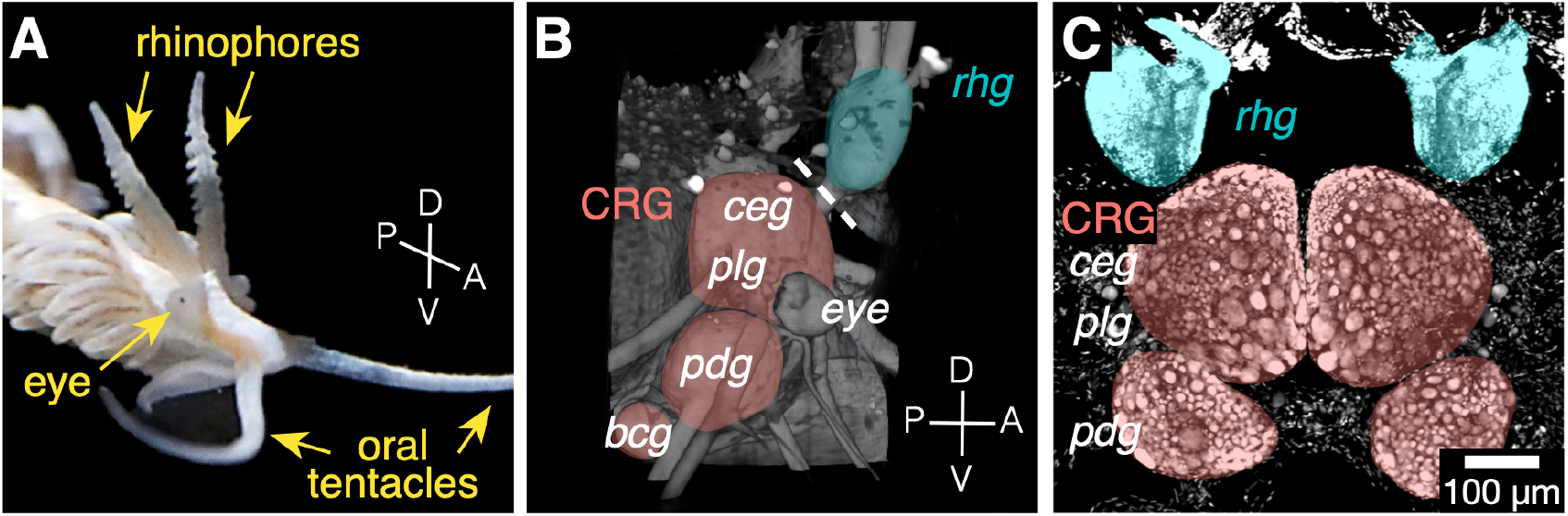
The *Berghia* brain atlas consists of cells taken from the central ring ganglia (CRG) and rhinophore ganglia (*rhg*). A) An adult *Berghia*, showing the head and positions of the rhinophores, eyes, and oral tentacles. B) 3D rendered and pseudocolored image of autofluorescence of the brain in an intact animal embedded in hydrogel. MHD-assisted CLARITY (***Dwyer et al., 2021***) was used to clear the tissue, then imaged using lightsheet microscopy. The right *rhg* (cyan) and CRG (pink) are shown. The dashed line crosses the *rhg* - *ceg* connective. C) Z-projection of a fluorescence confocal image of dissected head ganglia with DAPI-labeled nuclei. The CRG and *rhg* ganglia are labeled and colored corresponding to the 3D rendering in B. The *bcg* are not shown. Abbreviations: *ceg* - cerebral ganglion, *plg* - pleural ganglion, *pdg* - pedal ganglion, *bcg* - buccal ganglion.

Neuron type profiles that incorporate combinations of marker genes as well as soma size and position should enable the identification of neuronal sub-classes and even individual cell types in gastropods, as seen in well-studied laboratory species like *Caenorhabditis elegans* and *Drosophila melanogaster*.

*Berghia* offers many advantages over other commonly used gastropods for high throughput neuroscience research. It has a 2-month generation time, is commercially available, and can be reared and raised in the lab with minimal effort. Like other nudibranchs, the CRG, consisting of the fused cerebral (*ceg*) and pleural (*plg*) ganglia, pedal ganglion (*pdg*), and buccal ganglion (*bcg*), are condensed in the head (Fig. 1B, C). Homologs of neurons from other gastropods have already been found in *Berghia* (***Watkins, 2022***; ***Whitesel, 2021***).

Some key advantages of gastropod brains also present challenges for high-throughput scR-NAseq methods. First, there is a ten-fold variation in soma size, with the smallest neurons being less than 10*μ*m in diameter and the largest over 100*μ*m. Larger neurons are more fragile and less likely to survive cell dissociation, or may be too large to fit into the microfluidics devices of some scRNAseq methods. The CRG have fewer than 10,000 neurons overall, limiting the number of neurons that can be obtained from each sample. Finally, many neurons are present as a single bilateral pair per animal. Thus, obtaining the numbers of neurons typically used in scRNAseq studies is logistically challenging.

Besides the CRG, ganglia are associated with peripheral organs in gastropods, such as the genitals, tentacles, and rhinophores. In most nudibranchs, the rhinophore ganglion (*rhg*) is located distally within each rhinophore, a paired dorsal head appendage used for distance chemoreception (***Arey, 1918***; ***Cummins and Wyeth, 2014***; ***Storch and Welsch, 1969***; ***Wertz et al., 2006***). However, in *Berghia* the *rhg* sits at the base of the rhinophore and is separated from the CRG by a short nerve connective (Fig. 1B). Unlike the large neurons found in the CRG, no individual neurons or clusters have been identified in the *rhg*, due primarily to the small size of the neurons and their relative inaccessibility. Combining scRNAseq and HCR offers a new opportunity to catalog these peripheral neurons.

The results presented here describe neuronal gene expression in the CRG and *rhg*, showing that multiple neuronal classes, and non-neuronal cell types, can be recognized by differential gene expression. Other specific neuron types can be identified using the combination of gene expression with soma size and position. We found that unknown, unannotated genes represented a surprisingly sizable proportion of cluster-specific markers, highlighting importance of using unannotated genes for cell type identification, especially in understudied phyla like molluscs. We also present results showing unexpected diversity and organization of the neurons in the *rhg*, which suggest a high degree of complexity in this peripheral ganglion. Finally, we found differences in neuronal gene expression that distinguish the ganglia themselves and zones within ganglia. This study high-lights the use of modern, high-throughput molecular methods to develop a gene-based atlas of neurons in a non-traditional study species.

## Results

### We created a brain-enriched reference transcriptome for *Berghia*

Orthofinder2 (***Emms and Kelly, 2018***) was used to group similar sequences from transcriptomes, obtained from *Berghia* and other nudibranchs, as well as predicted peptides from the genomes of other gastropods like *Aplysia* and *Lottia*, into phylogenetically determined Hierarchical Orthogroups (HOGs). The output of the EnTAP (***Hart et al., 2020***) and Trinotate (***Bryant et al., 2017***) annotation pipelines on the *Berghia* transcriptome, as well as annotations from sequences from other gastropods were combined to annotate each HOG. The final reference transcriptome contained 78,000 transcripts representing 48,351 HOGs. Of the total number of HOGs, 22,844 contained sequences from both *Berghia* and at least one other gastropod, and the remaining 25,507 were found only in *Berghia*. Because of the gene-species tree reconciliation performed by Orthofinder2, some *Berghia* sequences were separated from their true orthologs into their own HOG, artificially inflating the number of *Berghia*-only HOGs. At least one annotation was appended onto 29,991 HOGs, and the remaining 18,360 HOGs were unannotated.

### Single cell transcriptomes

After gene counts for single cells were acquired using kallisto (***Bray et al., 2016***) and bustools (***Melsted et al., 2021***), data were analyzed using the R package Seurat (***Hao et al., 2021***). Following standard quality control filtering, we recovered transcriptomes for 872 cells from the CRG and 708 cells from the *rhg*. These 1,580 cells segregated into 19 clusters based on similarity of gene expression. Multiple clusters within the 19 were artifacts of overclustering our dataset, and so those not distinct in terms of differentially expressed genes (DEGs) were collapsed together, guided by a visualization of cluster stability across resolutions from Clustree (***Zappia and Oshlack, 2018***). This split-merge method allowed small clusters with strong signals, like the serotonergic neurons, to be separated out, instead of being subsumed into a larger cluster. A total of 14 clusters were established for the final dataset (Fig. 2).

**Figure 2.**
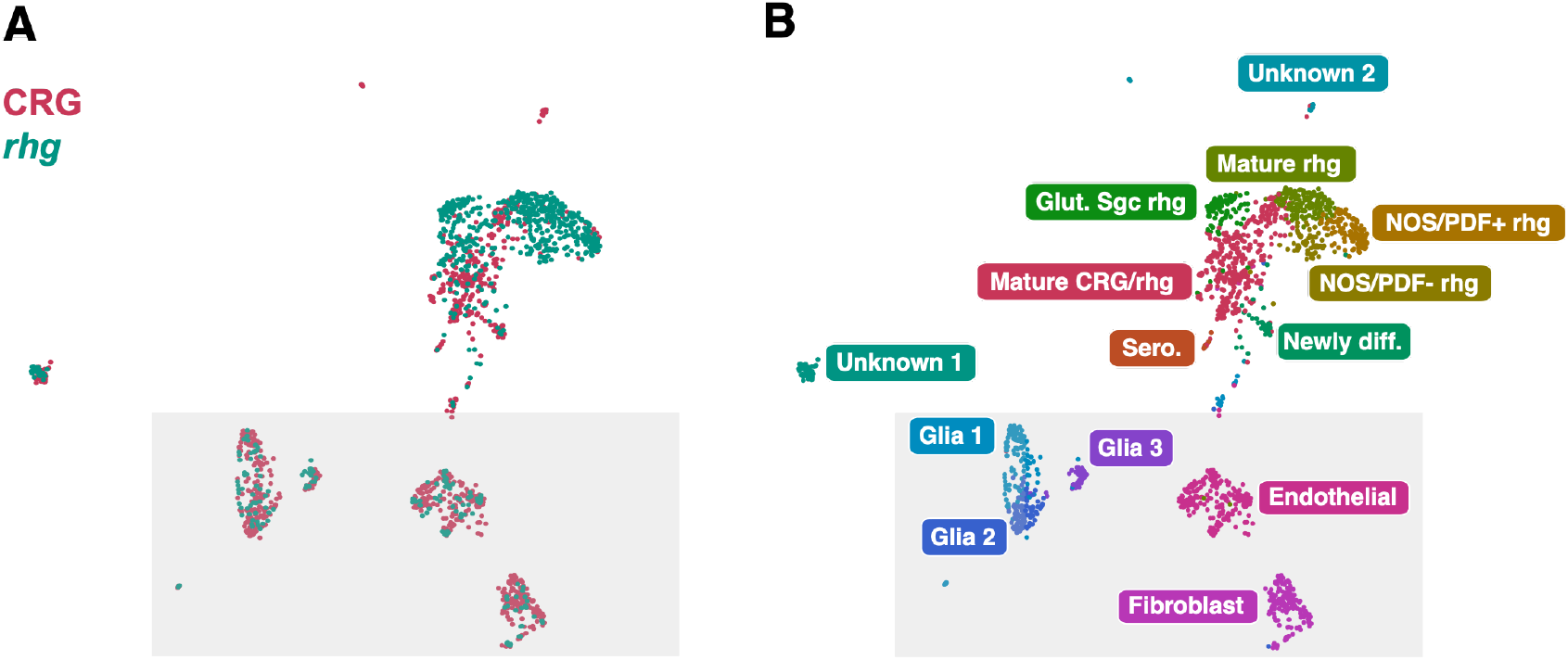
The *Berghia* single cell atlas contains 14 cell clusters, with 9 neuronal clusters and 5 non-neuronal clusters. A) Unifold Manifest Approximation and Projection (UMAP) plot of the *Berghia* single cell RNA-seq dataset from the two samples, the CRG (pink) and *rhg* (cyan). Most clusters are a mix from both samples. B) UMAP plot showing 1580 *Berghia* cells organized into 14 clusters. Non-neuronal clusters are contained within the shaded box.

Cell clusters were annotated using DEGs to assign putative identities based on the literature, and the sample origin. Nine neuronal and five non-neuronal cell clusters were found (Fig. 2B, Fig. 3), most of which had cells from both the *rhg* and CRG samples (Fig. 2A). Cluster markers were selected based on DEG between each cluster versus all others (Fig. 3). In most neuronal clusters, the best specific markers were often expressed in fewer than 50% of cells, indicative of heterogeneity within the clusters. This contrasted with the cluster markers for non-neuronal cell types, where the percentage of cells in the cluster expressing a marker was often much higher (over 90%). The remainder of our analysis focused on neuronal and glial clusters only.

**Figure 3.**
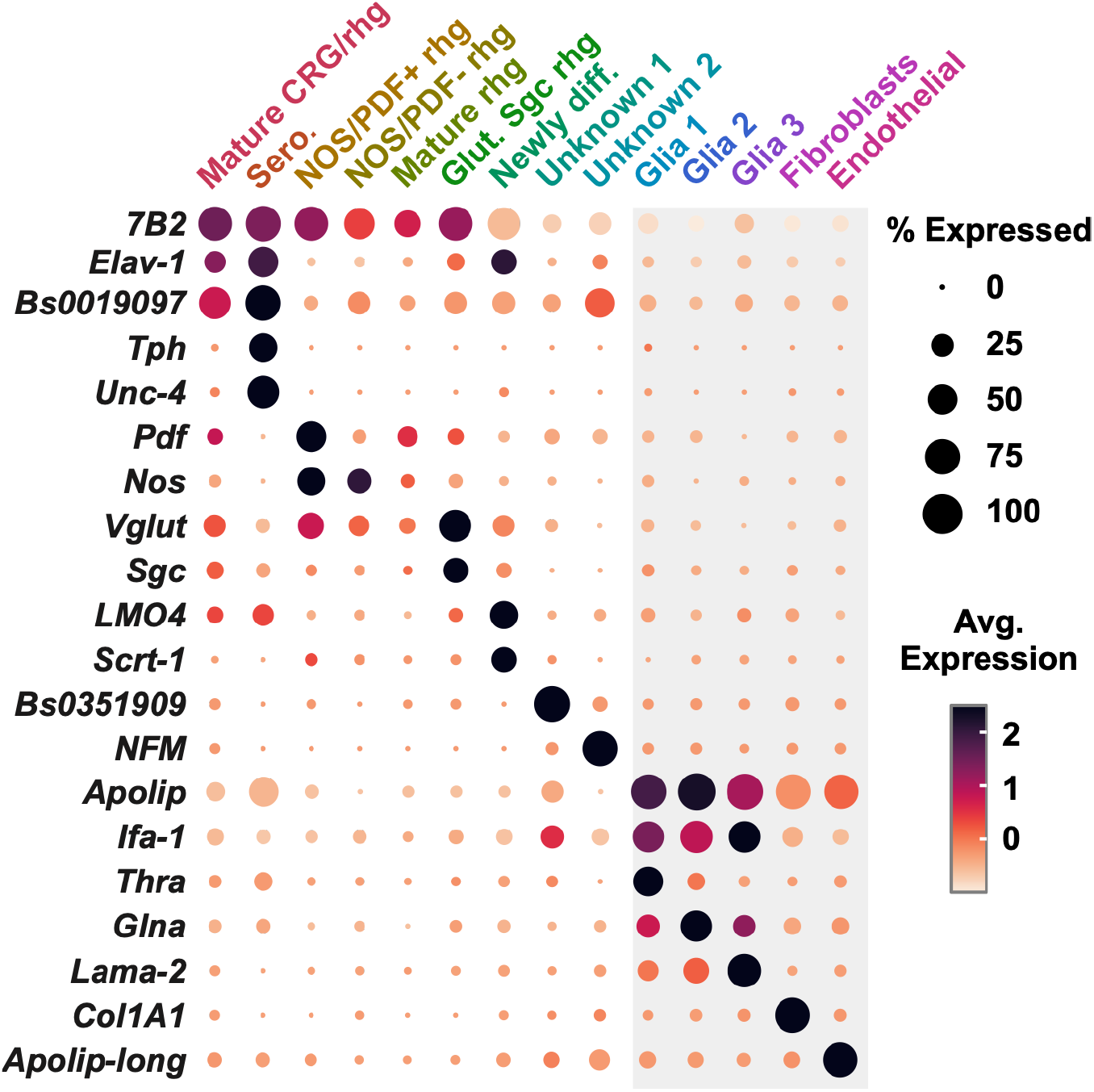
Dotplot of selected marker genes for each cluster. Non-neuronal cell types are boxed in gray. Cluster names and colors correspond to those in Fig. 2B.

### Pan-neuronal and pan-glial marker genes distinguish neurons and glia in the brain of *Berghia*

The expression of pan-neuronal genes marked neuronal clusters. 865 putative neurons were identified, spread across seven clusters (Fig. 3). Canonical neuronal markers like *Elav-1* (Fig. 3, Fig. 4A) were expressed primarily in four clusters. Oddly, *Elav-1* was not well represented in clusters primarily containing *rhg* neurons, where expression was mostly absent or sparse (Fig. 4A). Despite the low number of neurons with mRNA for *Elav-1* in the scRNAseq, HCR for *Elav-1* appeared to label all neurons, including neurons in the *rhg* (Fig. 4E), as expected from *Elav-1* expression in neurons in other animals.

**Figure 4.**
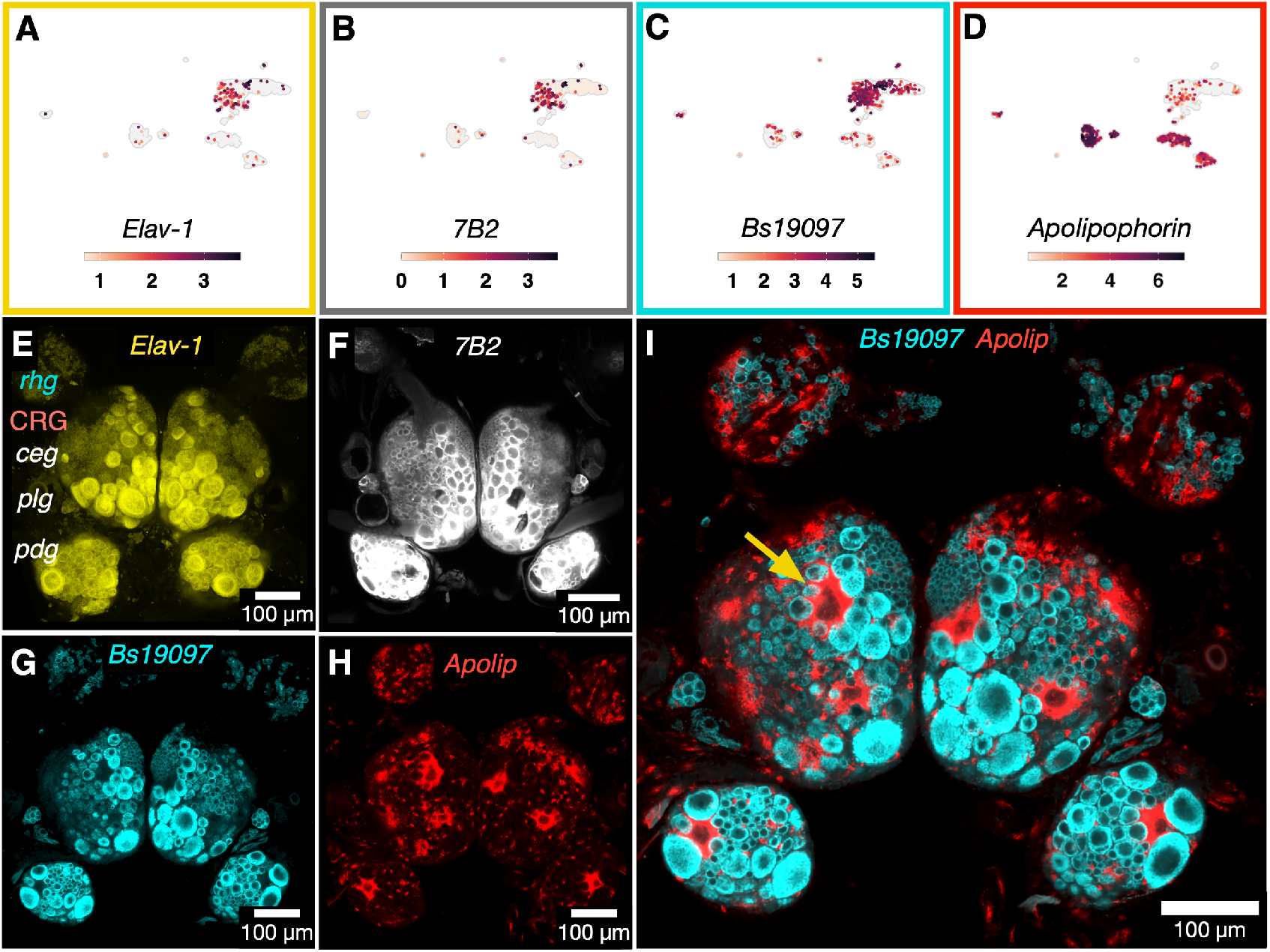
Pan-neuronal and pan-glial genes mark their respective cell types. A-D) UMAP plots showing mRNA abundances and distributions for *Elav-1* (A), *7B2* (B), *Bs19097* (C), and *Apolipophorin* (D). E-H) Single optical sections of a fluorescence confocal z-stack using HCR to label mRNA for pan-neuronal and pan-glial markers: *Elav-1* (E), *7B2* (F), unannotated transcript *Bs19097* (G), and the pan-glial marker *Apolipophorin* (H). Panels G & H are from the same multiplexed HCR sample. The yellow arrow indicates one of the individually identifiable giant glial cells. I) Expression of both *Bs19097* (cyan) and *Apolipophorin* (red) in the same sample.

Another pan-neuronal marker was neuroendocrine protein *7B2*. This gene is expressed in most neurons in other animals, including molluscs (***Hwang et al., 2000***; ***Marcinkiewicz et al., 1994***; ***Spijker et al., 1999***). *7B2* mRNA was abundant and found in all putative neuronal clusters regardless of sample origin (see Figs. 3, 4B & 4F). Low and sparse mRNA expression for *7B2* was also found in a cluster of putative non-neuronal glial and endothelial cells, consistent with previous findings in mouse (***Seidel et al., 1998***). *7B2* is the molecular chaperone for *prohormone/neuroendocrine convertase 2* (*pc2*), and *pc2* mRNA was also highly abundant in neuronal clusters. The expression of *7B2* and its molecular partner support the identification of cells in these clusters as neurons.

Finally, we found mRNA for an unannotated gene, *Bs19097*, present primarily in neuronal clusters and expressed ubiquitously in neurons across all ganglia (Fig. 3 & 4C). Multiplexed HCR showed extensive co-expression of *Elav-1* (Fig. 4E), *Bs19097* (Fig. 4G, I), and *7B2* (Fig. 4F), in addition to overlapping expression in presumptive neuronal clusters in the scRNAseq dataset (Fig. 4A-C). Because expression patterns for *Bs19097* closely match known pan-neuronal markers, we are confident in its identity as a pan-neuronal marker, despite its anonymity as a gene.

Three clusters containing putative glial cells shared many DEGs in common. Markers such as *glutamine synthetase* support the identity of cells in these clusters as glia (***Linser et al., 1997***) (Fig. 3). We did not find expression of canonical glial markers such as *Gfap* in vertebrates (***Eng, 1985***; ***Eng et al., 2000***) or *Repo* in arthropods (***Halter et al., 1995***; ***Xiong et al., 1994***) in the *Berghia* reference transcriptome, and therefore they were also absent from the scRNAseq data. This is consistent with recent scRNAseq in the brains of cephalopods molluscs (***Gavriouchkina et al., 2022***; ***Songco-Casey et al., 2022***; ***Styfhals et al., 2022***). A transcript annotated as an *intermediate filament protein 1* (*Ifa1*), the same superfamily as *Gfap* (***Peter and Stick, 2015***), was differentially expressed in the putative glial clusters (Fig. 3). *Apolipoprotein* (annotated as *Apolipophorin*) was used as a glial marker in *Octopus* (***Styfhals et al., 2022***). *Apolipohorin* in *Berghia* was differentially expressed in glial clusters, although mRNA for this transcript also appeared in cells from both endothelial and fibroblast clusters (Fig. 4D).

*Apoliophorin* HCR revealed a large number of small cells located in the sheath and neuropil that are consistent with glial positions and morphologies (Fig. 4H,I). Many small glial cells also appeared in between neuronal cell bodies. Additionally each ganglion contained multiple giant glia. These glia had giant nuclei, similar in size to some of the largest neuronal nuclei, and appeared to encase numerous small neuron somata (Fig. 4I, arrow). Giant glia were consistently found in bilaterally symmetrical pairs and were individually identifiable across animals. No cells were found expressing both neuronal and glial markers, supporting the specificity of markers for these broad cell types.

### The expression patterns of candidate genes and cluster markers subdivide neuronal classes

Candidate genes that subdivide neurons into large classes or groups were examined. These included markers for differentiating neurons, neurotransmitters, neuropeptides, and genes involved in transcriptional regulation. Most often, these markers spanned multiple neuronal clusters and by themselves did not distinguish clusters. Primary exceptions for this pattern were markers for newly differentiated neurons, which together cleanly delineated this cluster in the atlas. Serotonergic neurons also had highly specific cluster markers in the atlas. Overall, regardless of the specificity of a gene to a cluster in the single cell atlas, visualizing the expression in the brain revealed neuronal classes, with a wide diversity of neurons of different sizes, numbers, and locations within and between ganglia.

### A suite of transcription factors was expressed in newly differentiated neurons

Genes like *Lmo4*, *Sox6*, *Sox2*, *Scratch-1* are associated with neuronal differentiation. Only cells in only one cluster in the scRNAseq atlas expressed this suite of genes, suggesting that they were newly differentiated neurons (Fig. 5). *Lmo4* is a cofactor with *Neurogenin* for neural differentiation (***Asprer et al., 2011***). *Sox6* and *Sox2* work together to inhibit neural differentiation in vertebrates (***Lee et al., 2014***; ***Li et al., 2022***). *SoxB1* (homolog of vertebrate *Sox* family genes) is key to neuroblast formation in *Drosophila* (***Buescher et al., 2002***). *Scratch-1* is important for neural differentiation in *C. elegans*, mammalian cell culture, and *Drosophila* (***Manzanares et al., 2001***; ***Nakakura et al., 2001***; ***Nieto, 2002***). In mammalian forebrain, *Scratch* genes act downstream of proneural genes like *Neurogenin* and *Ascl1* and control the initiation of migration of newly differentiated neurons (***Itoh et al., 2013***).

**Figure 5.**
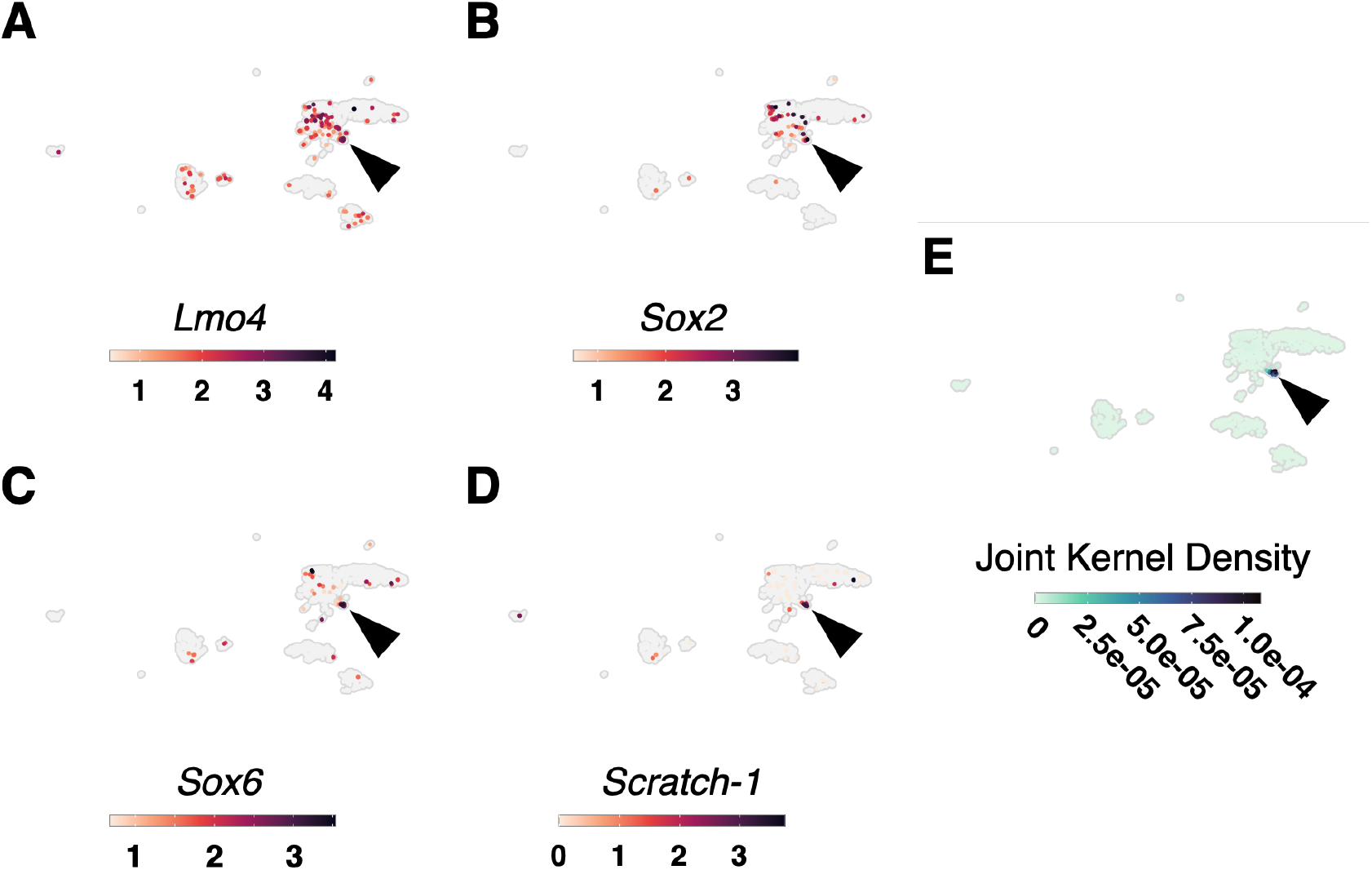
Marker genes expressed in a cluster containing newly differentiated neurons. A-D) *Lmo4* (A), *Sox2* (B), *Sox6* (C), and *Scratch-1* (D). E) Joint kernel density plot showing the peak of overlapping expression of the four genes specifically in the newly differentiated neuron cluster (arrowheads).

### Major small molecule neurotransmitters: glutamate, GABA, acetylcholine and serotonin

We looked for the expression of candidate enzymes and transporters associated with small molecule neurotransmitters. In the single cell atlas, mRNAs were found for *Vesicular glutamate transporter* (*Vglut*, glutamate), *Choline acetyltransferase* (*Chat*, acetylcholine), *Glutamate decarboxylase* (*Gad*, GABA), *Tryptophan hydroxylase* (*Tph*, serotonin), *Histidine decarboxylase* (histamine), and *Tyramine beta-hydroxylase* (octopamine). There was no evidence of mRNA for *Tyrosine hydroxylase*, the rate-limiting enzyme for catecholamine synthesis, but this may be the result of low gene expression, the small number of neurons captured, and the rarity of catecholaminergic neurons in the CRG of gastropods (***Croll, 2001***) rather than true absence. A total of 484 neurons (about 50%) possessed mRNAs for at least one of these genes (Fig. 6A-D).

**Figure 6.**
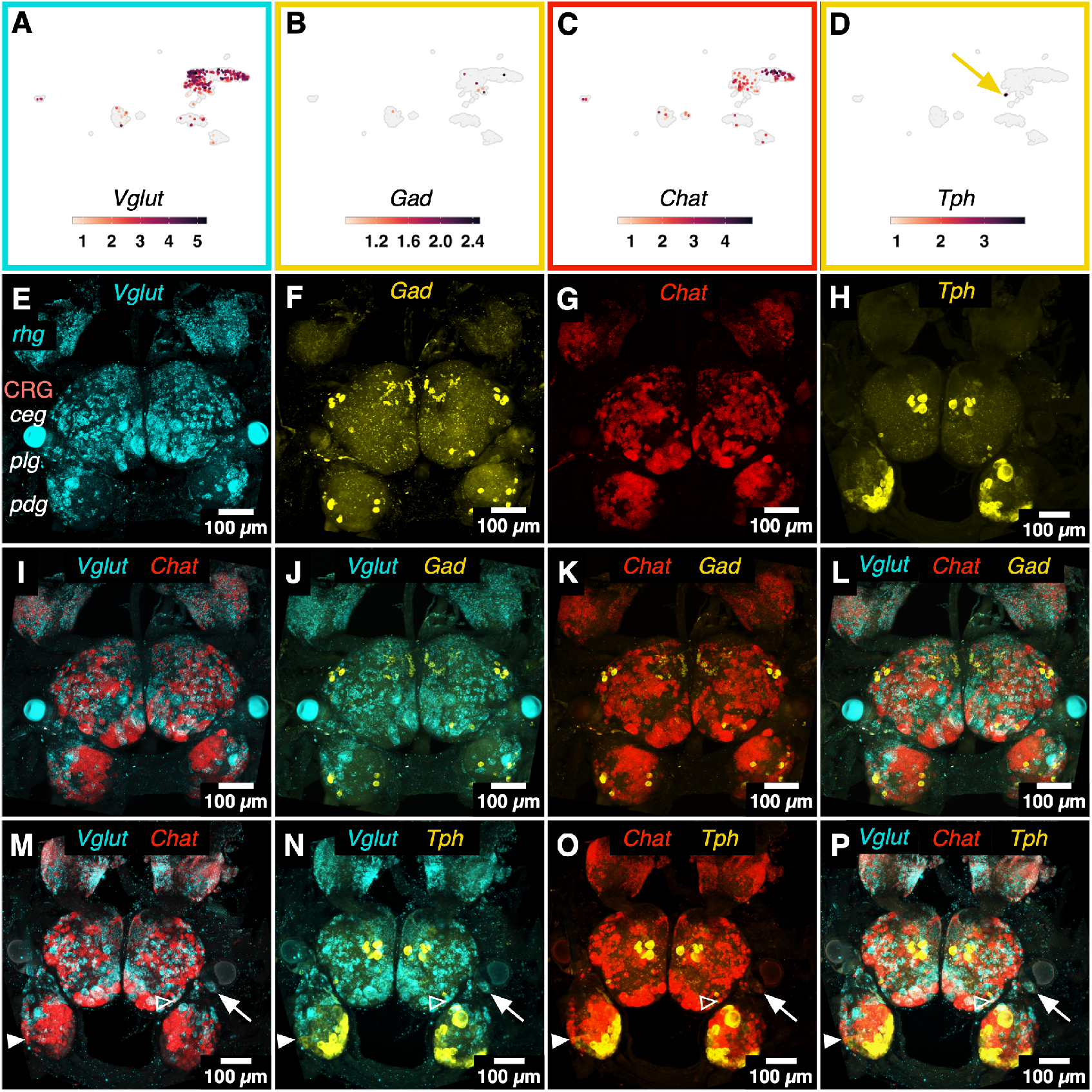
Abundance and visualization of mRNA for neurotransmitter-associated enzymes and transporters in *Berghia*. A-D) UMAP plots showing mRNA abundances of *Vglut* (A), *Gad* (B), *Chat* (C), and *Tph* (D) across neuronal clusters. The yellow arrow in (D) points to the small cluster of *Tph*+ neurons. E-H) Z-projections of fluorescence confocal images using HCR to label *Vglut* (E), *Gad* (F), *Chat* (G), and *Tph* (H). (E-G) are channels from the same multiplexed sample; (I-L) show *Vglut*, *Gad*, and *Chat* pairwise and triple-labeled from that same sample. (M-P) show *Vglut*, *Chat* and *Tph* pairwise and triple-labeled in the sample shown in (H). The white arrows point to a *Vglut*+ photoreceptor in the eye. The open white arrowheads point to a neuron double-labeled for *Vglut* and *Chat*. The solid white arrowheads point to a neuron double-labeled for *Chat* and *Tph*. Most of the neurons are labeled for only one of the genes. **Figure 6–Figure supplement 1.** Co-expression of neurotransmitter-related genes in single cells

Using HCR, we visualized mRNAs for the most prevalent neurotransmitter-associated genes: *Vglut*, *Gad*, *Chat*, and *Tph* (see Fig. 6E-P). A large percentage of the neurons in the head ganglia expressed the genes associated with these neurotransmitters. There was co-expression of at least two neurotransmitters in a small number of neurons for *Vglut*, *Gad*, *Chat*, and *Tph* (Fig. 6M-P, arrowheads).

Glutamate appeared to be the most prevalent neurotransmitter in the head ganglia based on number of cells containing mRNA for *Vglut*. *Vglut* loads glutamate into vesicles to be released at synapses. One *Vglut* transcript was expressed in 327 neurons from both the CRG and *rhg* scRNAseq samples (Fig. 6A). The spatial distribution of *Vglut* mRNA in the brain supports the single cell data– *Vglut* HCR labeled many neurons and at relatively high levels as a transporter (Fig 6E). *Vglut* HCR also labeled photoreceptors in the eye (Fig. 6M-P, arrow).

Based on the expression of its rate-limiting synthesis enzyme, *Gad*, GABA (*γ-aminobutyric acid*) appears to not be a prominent neurotransmitter in the brain of *Berghia*; only a handful of neurons containing *Gad* mRNA were found in the single cell dataset (Fig. 6B). *Gad* HCR labeled only a small number of neurons across the CRG (Fig. 6F), which is similar to what was seen for GABA IHC (***Gunaratne et al., 2014***; ***Gunaratne and Katz, 2016***). However, the mRNA expression levels in many of these neurons were higher that of *Vglut* and *Chat* based on the density and intensity of the HCR signal in *Gad* neurons. There were no clearly distinguishable *rhg* neurons that expressed *Gad*.

Acetylcholine, like glutamate, appeared widespread in neurons throughout the head ganglia. *Chat*, the rate-limiting enzyme for acetylcholine synthesis, was expressed in multiple clusters of mature neurons from the CRG and *rhg* scRNAseq dataset (Fig. 6C). 108 neurons contained *Chat* mRNA in the scRNAseq dataset. *Chat* HCR showed mRNA present in many neurons (Fig. 6G), but fewer than those that expressed *Vglut* (Fig. 6E), which is consistent with the percentage found in the brain of the slug Limax (***D’Este et al., 2011***).

Serotonin was not as widespread as glutamate or acetylcholine but was more prevelant than GABA. mRNA for *Tph*, the rate limiting enzyme for serotonin synthesis, was found exclusively in a cluster containing only 8 neurons in the single cell atlas (Fig. 6D, yellow arrow). However, *Tph* HCR showed mRNA in the brain was more prevalent than suggested from the single cell data (Fig. 6H). Unlike *Vglut* and *Chat*, HCR for *Tph* was much more spatially restricted. The majority of *Tph*- expressing cells were found in the pedal ganglia, and a much smaller number (< 20) were found in the cerebral-pleural ganglia (Fig. 6H). The distribution of *Tph* neurons in the *Berghia* brain matches the highly conserved pattern of serotonin IHC seen in other nudibranch species (***Newcomb et al., 2006***).

Co-expression of multiple genes associated with different neurotransmitters (***Hnasko and Edwards, 2012***) was found in both the scRNA-seq dataset and with HCR. The small molecule neurotransmitters were mostly non-overlapping in expression (Fig. 6I-P). However, at least one neuron in the *plg* colabeled for *Vglut* and *Chat* (Fig 6 M, P open arrowhead) and another in the *pdg* expresses both *Tph* and *Chat* (Fig. 6 O,P, solid arrowhead).

### Conserved transcriptional regulators and signaling pathway genes define brain regions and neuronal classes

*Six3/6* is involved in many different developmental processes in animals, including anterior brain and eye development (***Bernier et al., 2000***; ***Seimiya and Gehring, 2000***). While *Six3/6* mRNA was found in neurons in multiple clusters in the atlas, it was highly differentially expressed in the glutamatergic *Solute guanylate cyclase* (Sgc) *rhg* neuronal cluster (Fig. 7A). Using HCR to visualize *Six3/6* mRNA, expression was distributed among many *rhg* neurons while in the CRG it was essentially restricted to *ceg* neurons (Fig. 7C). These are the most anterior ganglia of the major head ganglia (see Fig. 1B). Anterior expression is consistent with *Six6* expression in developing vertebrate brains (***Oliver et al., 1995***). *Six3/6* also has a role in eye development in multiple animal phyla (***Seimiya and Gehring, 2000***; ***Seo et al., 1998***), but using HCR, there was no expression of *Six3/6* in the eye. However, a single neuron within the optic ganglia was labeled for *Six3/6*. *Optix* (the *Six3/6* homolog in *Drosophila*) regulates and demarcates the larval optic lobe (***Gold and Brand, 2014***).

**Figure 7.**
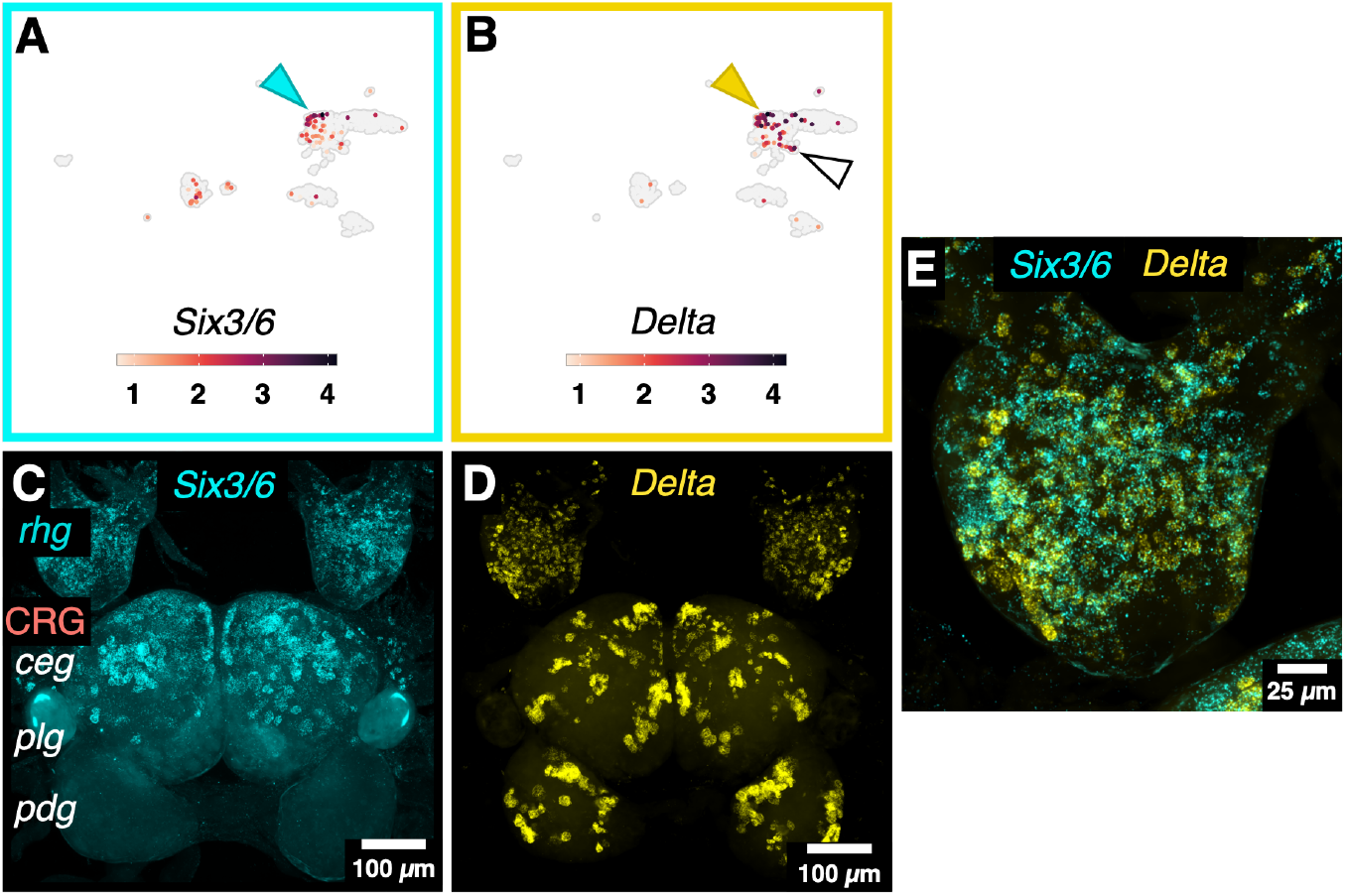
Expression of transcriptional regulators and signaling pathway genes help define different brain regions. A) *Six3/6* mRNA was most abundant in *Soluble guanylate cyclase* (*Sgc*) *rhg* neurons in the single cell atlas and was differentially expressed in this cluster (cyan arrowhead). B) *Delta* mRNA was concentrated in the *Sgc rhg* (yellow arrowhead) and differentiating neuron (white arrowhead) clusters in the single cell atlas. Delta was differentially expressed in the *Sgc rhg* neuron cluster. C) Z-projection of a fluorescence confocal image using HCR to label *Six3/6* mRNA. It was almost exclusively present in neurons in the anterior-most ganglia, the *rhg* and the cerebral ganglion *ceg*. D) Z-projection of a fluorescence confocal image of the left *rhg* using HCR to label *Delta* mRNA showed it was widely distributed across the ganglion. Many neurons in the *rhg* contained *Delta* mRNA, some of which likely correspond with the *Sgc rhg* neurons. E) Higher magnification of the *rhg* showing co-expression of *Six3/6* and *Delta* in many of the same neurons. C, D, and E are the same sample.

A transcript annotated as *Delta-like* was another broadly expressed gene in the single cell data. Both *Delta* and *Jagged* are similar transmembrane protein ligands for Notch receptors to initiate a signaling cascade typically associated with early development of major aspects of nervous systems across animals (***Bettenhausen et al., 1995***; ***Chitnis et al., 1995***; ***Henrique et al., 1995***; ***Kawaguchi et al., 2008***). In *Berghia*, there was a higher density of *Delta*-expressing neurons in the *rhg*, and *Delta* was a DEG in *rhg* neurons (Fig. 7B). *Six3/6* and *Delta* were DEGs in the glutamatergic *Sgc rhg* neuron cluster in the scRNAseq atlas (Fig. 7B,D). Many neurons in the *rhg* expressed both *Six3/6* and *Delta* (Fig. 7E). *Delta* mRNA was present in the scRNAseq cluster of newly differentiated neurons, consistent with its role in development. However, *Delta* mRNA was also found in neurons across all head ganglia in presumably mature neurons from adult CRG (Fig. 7D). Many of the *Delta* expressing neurons differ in size and location, suggesting that *Delta* may be part of the gene expression profiles of multiple neuron types in the *Berghia* CNS (Fig. 7D).

### The neuropeptide complement in *Berghia* includes both broadly conserved bilaterian genes and others that are lineage-specific

At least 40 neuropeptides were found in the scRNAseq data, representing approximately 30 families out of a minimum set of 65 molluscan neuropeptides (***De Oliveira et al., 2019***) (Fig. 8). The large Mature CRG/*rhg* neuron cluster showed high average expression of most neuropeptides in a small number of neurons, suggesting this cluster consists of many rare neuron types. *APGWamide* (*Apgw* or *Cerebral peptide*) was one of the most highly expressed genes overall. Visualization of *Apgw* mRNA in the CRG and *rhg* reflected both the wide range of cells expressing this neuropeptide and the extremely high levels of expression (Fig. 9A); *Apgw* mRNA was present in both the somata and axons of many neurons. *CCWamide* was differentially expressed in the largest neuronal clusters, and widely expressed across clusters in the scRNAseq dataset (Fig. 8, 9B). *CCWamide* was recently identified in a broad search for neuropeptides in annelids, molluscs, and other lophotrochozoan phyla (***Thiel et al., 2021***; ***Williams et al., 2017***). Its expression pattern in molluscan brains was not previously known. HCR for *CCWamide* in *Berghia* reflected the high abundance and broad expression of *CCWamide* seen in the single cell atlas (Fig. 9B).

**Figure 8.**
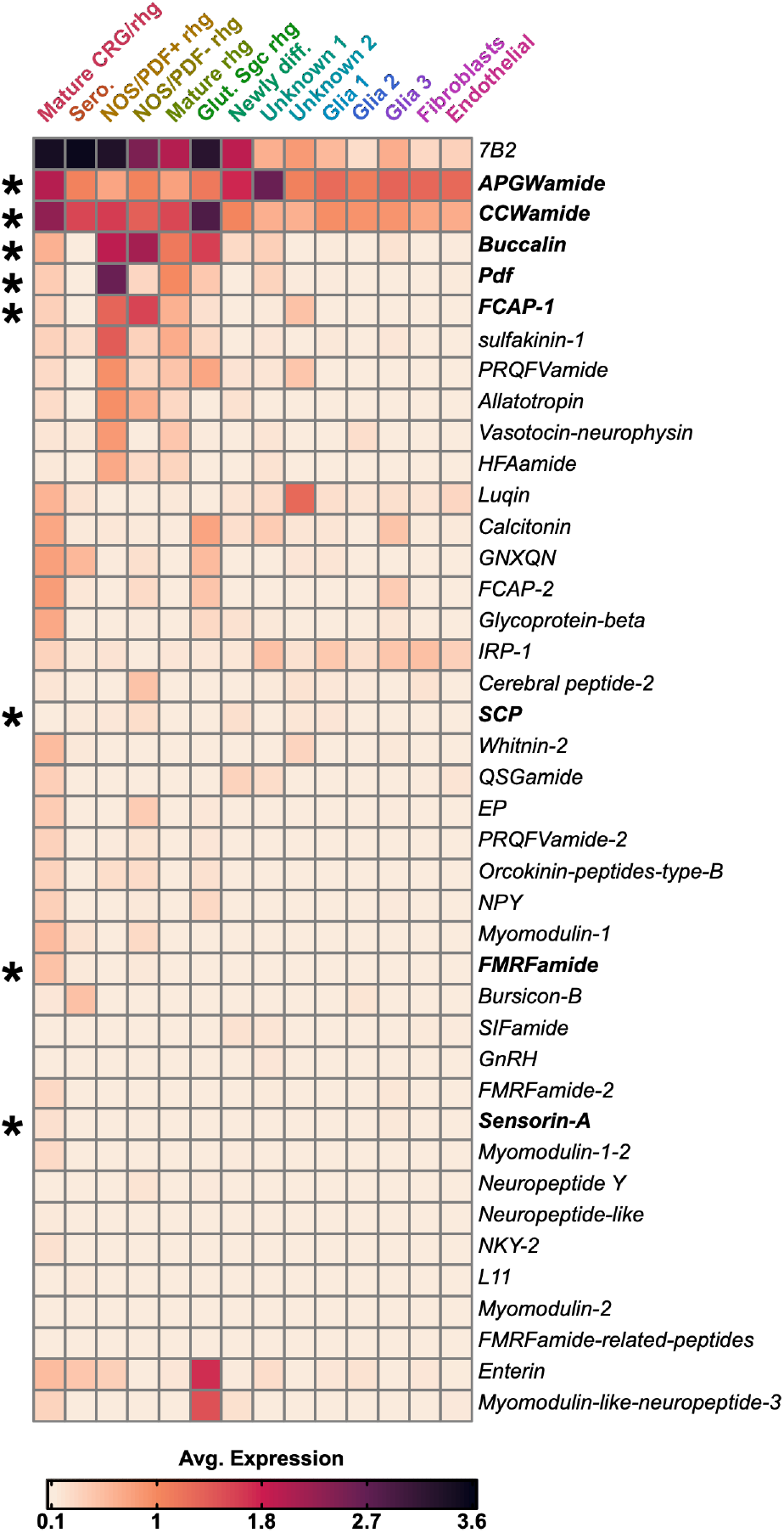
Over 40 types of neuropeptides were found within the atlas, and their expression varied by cluster. The mature CRG/*rhg* neuron cluster contained the largest number of neurons, and showed the largest and most varied expression of neuropeptides between the clusters. Only a few neuropeptides were restricted to specific clusters. *Luqin* was primarily expressed in the Unknown 1 cluster of neurons. *Enterin* and *Myomodulin-like-neuropeptide-3* expression marked neurons in the glutamatergic, *Sgc+ rhg* cluster. Asterisks indicate neuropeptides that were selected for visualization using HCR in Figs 9 & 10.

**Figure 9.**
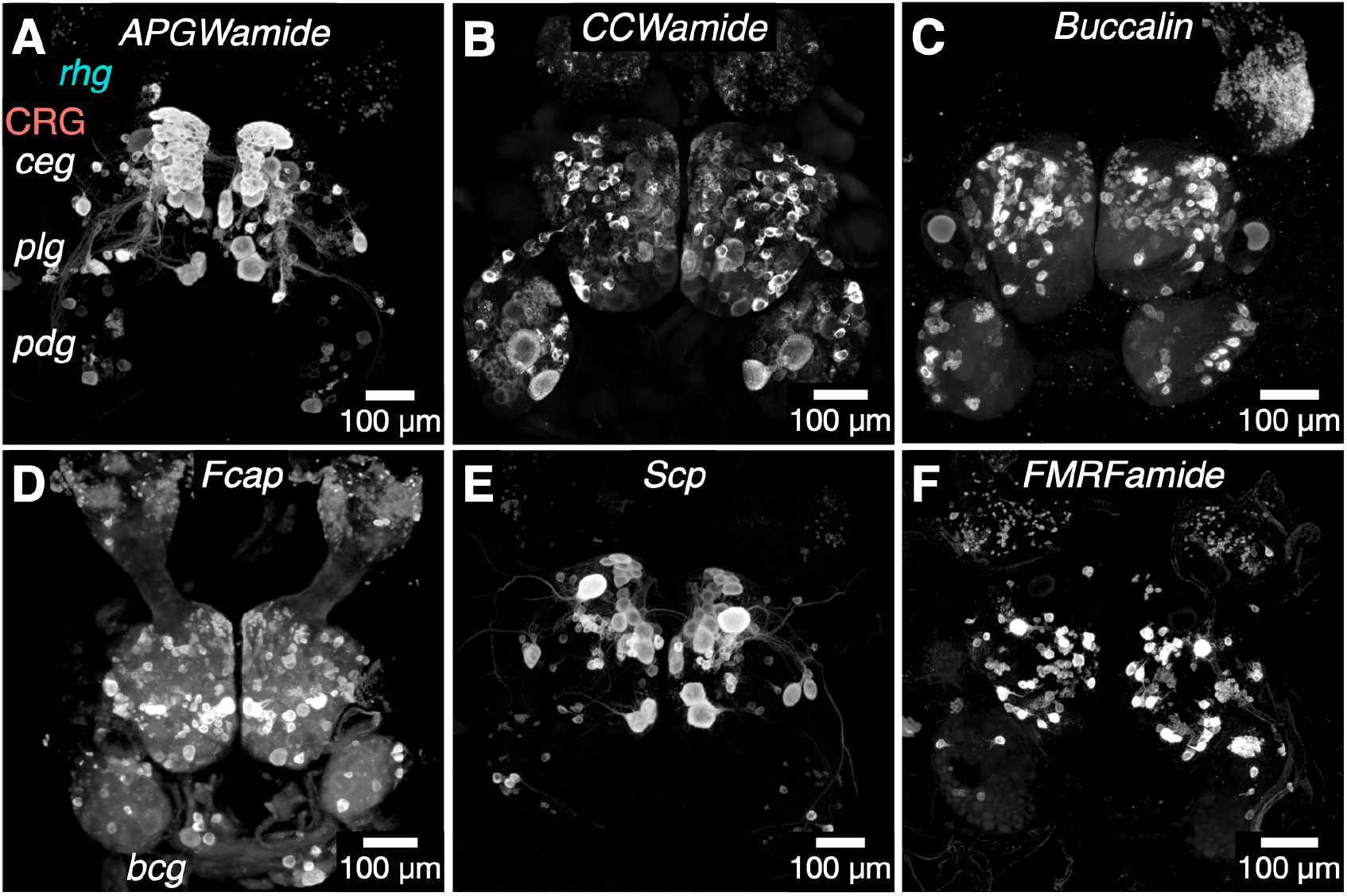
Neuropeptide expression was distinct for each gene, and varied greatly in the number of neurons and their distributions across the ganglia. Z-projections of a fluorescence confocal image using HCR to label mRNA for eight neuropeptides: *APGWamide* (A), *CCWamide* (B), *Buccalin* (C), *Fcap* (D), *Scp* (E), and *FMRFamide* (F).

Other neuropeptides showed more restricted expression. Except for some neurons in the mature CRG/*rhg* cluster, *Luqin* is found only in the Unknown 2 cluster of neurons, and *Irp-1* in the Glial 3 cluster. In the *Nos/Pdf+ rhg* cluster, *Pigment-dispersing factor* (*Pdf*, also called *Cerebrin*, Fig. 10H) and one transcript of *Feeding circuit-activating peptide* (*Fcap*, Fig. 9D) were DEGs. Yet mRNA for these peptides were not restricted to the *Nos/Pdf+ rhg* neuron cluster, was present in other neuronal clusters, and was spatially distributed in many, if not all, ganglia. For example, HCR shows *Pdf* mRNA is also highly expressed by a few distinct CRG neuron types in the *ceg* (Fig. 10H). Besides its expression in small *rhg* neurons, *Fcap* is also expressed in some of the largest neurons within the lateral region of the *rhg*, as well as many of the large neurons of the *ceg* and *plg* (Fig. 9E). One large and 4-5 other smaller *bcg* neurons also express *Fcap* (Fig. 9D). *Buccalin*, a putative homolog for *Allatostatin-A* (***Veenstra, 2010***), was also differentially expressed in the *Nos/Pdf+ rhg* cluster (Fig. 9C). Numerous *Buccalin*-expressing cells were found primarily in the *rhg*, *ceg*, and *pdg*, but only a handful were present in the *plg* (Fig. 9D).

**Figure 10.**
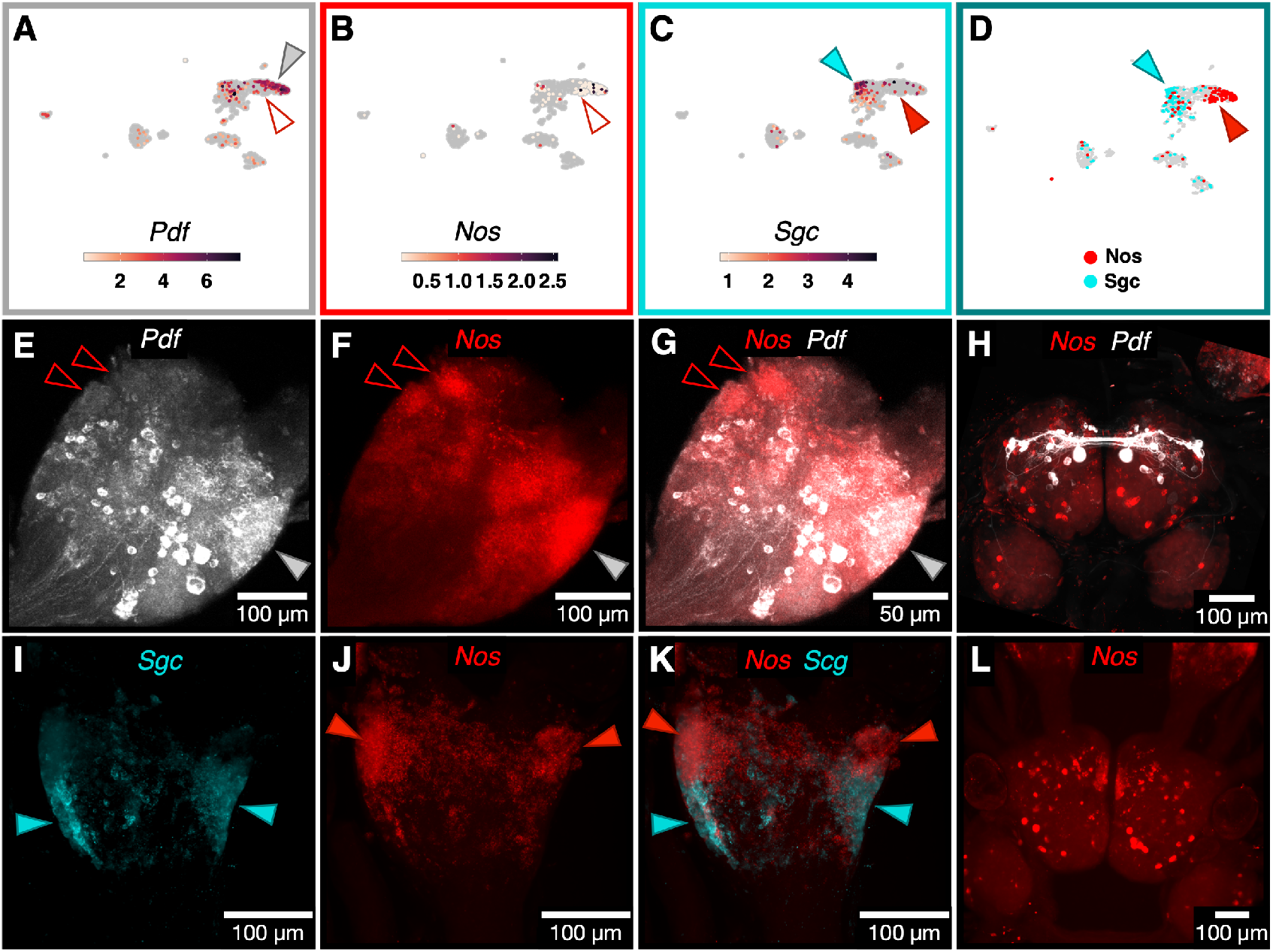
Differences in location and co-expression within *Nos*-expressing neuronal populations are matched by spatial segregation. A-D) UMAP plots of mRNA abundances of the neuropeptide *Pdf* (A), *Nos* (B), and *Sgc* (C) in the single cell atlas. D) UMAP plots showing mutually exclusive expression of either *Nos* (red), or *Sgc* (cyan) in some *rhg* neuron populations. E-G) Z-projections of a fluorescence confocal image using HCR to label mRNA for *Pdf* (E), *Nos* (F), and the two channels merged (G) in the *rhg*. Open red arrowheads show *Nos*-expressing cells, and the closed arrowhead shows *Pdf* -expressing cells. H) Z-projection of a fluorescence confocal image using HCR to label mRNA for *Pdf* (white), *Nos* (red) in the CRG. A pair of medium sized *Pdf+* neurons sit near the midline in the *ceg* and send projections contralaterally. The projections appear to meet other *Pdf+* neurons in the *ceg*, which together form a distinct loop throughout the ganglia. I-K) Z-projections of a fluorescence confocal image using HCR to label mRNA in the rhinophore ganglia for *Sgc* (I), *Nos* (J) and the merged image (K). There were distinct populations of cells expressing each gene in the *rhg*. Closed red arrowheads indicate *Nos*+ neuron populations, and closed cyan arrowheads indicate *Sgc*+ neuron populations. L) Z-projection of fluorscence confocal stack using HCR to label mRNA for *Nos* (red) in the CRG.

In addition to investigating neuropeptides that were differentially expressed in the single cell atlas, we also looked for expression of neuropeptide candidates gleaned from the literature, including *Small cardioactive peptide* (*Scp*) (Fig. 9E) and *FMRFamide* (Fig. 9F). Like the differentially expressed neuropeptides, these candidate neuropeptides were present in neurons of many different positions and size classes within the CRG and *rhg*.

### Many *rhg* neurons are distinguished by genes involved in Nitric oxide (NO) signaling

Two clusters of *rhg* neurons shared expression of *Chat* and *Nitric oxide synthase* (*Nos*), but were distinct in the presence of *Pigment-dispersing factor* (*Pdf*) mRNA (Fig. 10A,B). HCR for *Nos* and *Pdf* showed two populations of neurons that likely corresponded to the *Nos/Pdf+* and *Nos/Pdf-* cells in the single cell dataset (Fig. 10E-G). Both *Pdf* and *Nos* mRNA were also present in a small number of CRG neurons in the scRNAseq data and labeled using HCR (Fig. 10H).

The other primarily *rhg* neuron cluster in the scRNAseq atlas was marked by the expression of *Vglut* and *Soluble guanylate cyclase* (*Sgc*). *Sgc* is the receptor for nitric oxide (NO) (***Martin et al., 2005***), raising the possibility that these neurons may receive NO as signals, potentially from other *rhg* neurons that express *Nos*. When co-labeled with *Nos* using HCR in the *rhg*, mRNA for the two genes were mostly mutually exclusive (Fig. 10D,I-K). There appeared to be fewer, larger *rhg* neurons that express *Sgc*, which were interspersed among *Nos* expressing neurons (Fig. 10K).

### Specific neuronal cell classes and types are distinguishable based on their molecular signatures

Some *Berghia* neurons express the bilaterian molecular signature of mechanosensory neurons. Across bilaterians, mechanosensors, as well as interneurons that synapse with mechanosensors, share a similar molecular profile. They are glutamatergic, based on their expression of *Vesicular glutamate transporter* (*Vglut*), and express the transcription factors *Brain-3* (*Brn3*), *Dorsal root ganglion homeobox* (*Drgx*), *Islet-1* (*Isl1*) and *Lim-homeobox 3/4* (*Lhx3/4*) (***Nomaksteinsky et al., 2013***). *Brn3* and *Vglut* HCR localized mRNA to neurons in the *plg*, consistent with the known mechano and nociceptive circuits in *Aplysia* (***Walters et al., 2004***). The neuropeptide *Sensorin-A* (*SenA*) was also expressed along with *Brn3+* mechanosensory neurons in *Lymnaea* (***Nomaksteinsky et al., 2013***). However, most *Brn3+* neurons in *Berghia*’s scRNAseq dataset were not *SenA+*. A small number of neurons within the larger mature CRG/*rhg* atlas cluster were both *SenA+* and *Brn3+* (Fig. 11). Consistent with co-expression results from the atlas, HCR for these genes showed co-expression in only a few neurons (Fig. 11G, dotted box). S-cell homologs (***Getting, 1976***) in *Berghia* consisted of about 15 small *SenA+* neurons. These cells did not co-express *Brn3*, but there were 1-2 somewhat larger *Brn3+* neurons nestled among them. *Berghia* S-cells were glutamatergic, based on their expression of *Vglut*, seen both in the atlas and using HCR. S-cells in other nudibranchs are also glutamatergic (***Megalou et al., 2009***).

**Figure 11.**
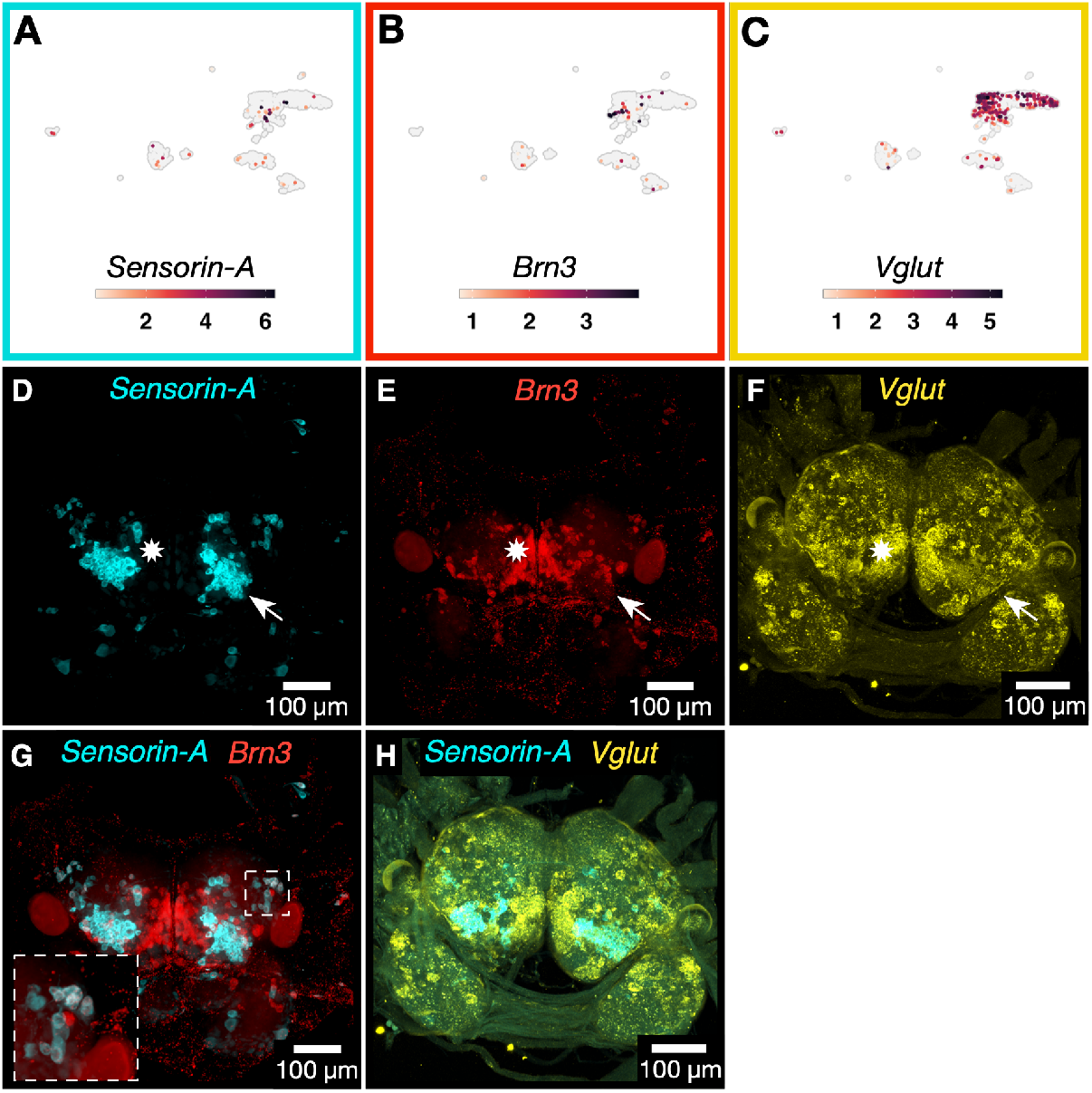
Co-expression of *Vglut* in *Brn3+* cells indicates mechanosensory neuronal identity. A-C) UMAP plots showing expression of *Sensorin-A* (A), *Brn3* (B) and *Vglut* (C) in the single cell dataset. Z-projection of a fluorescence confocal image using multiplexed HCR to label mRNA for D) *Sensorin-A*, and E) *Brn3* in the same sample. F) Z-projection of a fluorescence confocal image HCR to label mRNA for *Vglut*. White arrow indicates the likely homologs of the S-cells (***Getting, 1976***) known from other nudibranchs, expressing both *SenA* and *Vglut*, but not *Brn3*. White star indicates cell populations that express *Brn3* and *Vglut*, but not *SenA*. G) A merged z-projection of fluorescence confocal images of multiplexed HCR for *SenA* and *Brn3* in the same sample as (D,E). Dotted box indicates closeup of *Sen-A*+/*Brn3*+ neurons in the *ceg*. H) A merged z-projection of fluorescence confocal images of multiplexed HCR for *SenA* and *Vglut* in the same sample as (F).

### Transcription factor *Unc-4* associates with serotonergic efferent neurons in *Berghia*

*Unc-4* underlies specification of motor neurons in both *C. elegans* and *Drosophila* (***Lacin et al., 2020***; ***Pflugrad et al., 1997***). Motor neurons in these species are also cholinergic and express *Chat*, and *Unc-4* is known in *Drosophila* to repress a GABAergic cell fate leading to a cholinergic one (***Lacin et al., 2020***). Surprisingly, in the *Berghia* single cell atlas, *Unc-4* and *Solute carrier family 46 member 3* (*Scf46m3*) were differentially expressed along with *Tryptophan hydroxylase* (*Tph*). Here, *Tph* mRNA was restricted to a small cluster (8 cells), while *Unc-4* and *Scf46m3* mRNAs were found more widely (Fig. 12). Consistent with the co-expression seen in the single cell data, *Unc-4* and *Scf46m3* were co-expressed in CRG neurons along with *Tph*, though both genes were also expressed in a small number of neurons that did not express *Tph* (Fig. 12A-G, cyan and red arrowheads). As the rate-limiting step in serotonin production, *Tph* was expected to be found only in serotonergic neurons. Further supporting the specificity of *Unc-4* and *Scf46m3* to serotonergic, rather than cholinergic, neurons was the lack of overlapping *Chat* and *Tph* mRNA in the atlas. Visualizing mRNAs for *Chat* and *Tph* with HCR showed overlap in only a handful of neurons (see Fig. 6O,P).

**Figure 12.**
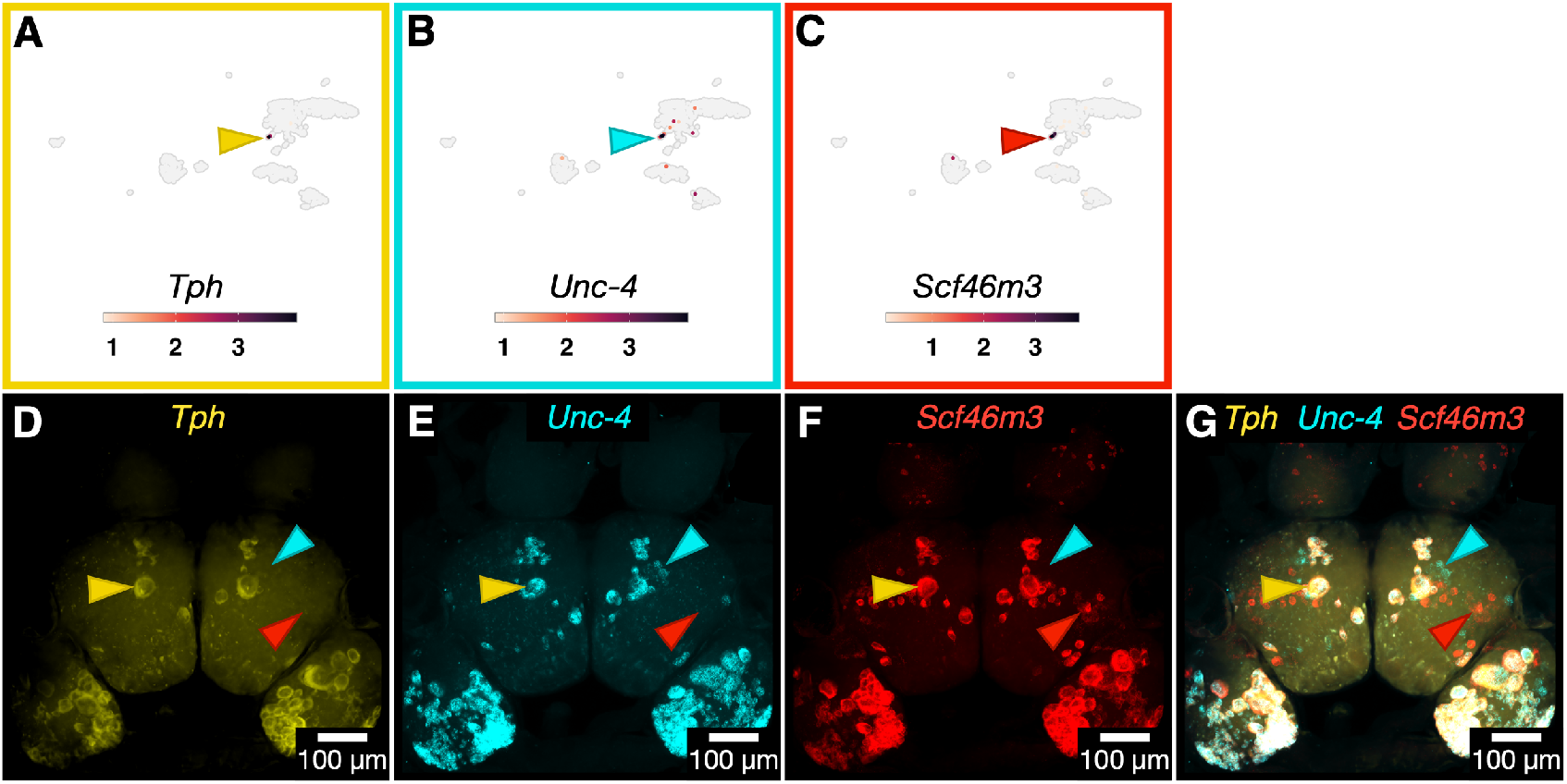
HCR labeling shows mRNA for *Tryptophan hydroxylase* (*Tph*), *Unc-4* and *Solute carrier family 46 member 3* (*Scf46m3*) were co-expressed in serotonergic neurons. A-C) UMAP plots showing mRNA abundances for *Tph* (A), *Unc-4* (B), and *Scf46m3* (C). Arrowheads point to the cluster where all three genes are found. D-G) Z-projection of a fluorescence confocal image using HCR to label *Tph* (D), *Unc-4* (E), and *Scf46m3* (F). G) The merged image showing the overlap of these 3 genes. The yellow arrowheads point to the likely homolog of the *Aplysia* metacerebral cell, seen in all three channels. The cyan arrowhead point to an example of neurons expressing only *Unc-4*. The red arrowhead point to an example of neurons expressing only *Scf46m3*.

### Detection of molecular signatures for known identifiable neurons

A visually identifiable bilaterally symmetric neuron sits on the ventral surface of the *ceg*, near the anterior-most region of the ganglion. It is surrounded by a field of much smaller, equally sized neurons and is the largest of neurons in the ventral *ceg* (Fig. 13A). Due to its unique size and location alone, it was straightforward to identify this neuron in images of the ventral surface of the CRG using only a nuclear label. This neuron was seen in images for pan-neuronal genes, as expected, but also multiple neuropeptides, including *APGWamide*, *Scp*, and *Fcap* (Fig. 13B-D). Besides neuropeptides, the neuron was observed in HCR images for the enzyme *Chat* (Fig. 13E).

**Figure 13.**
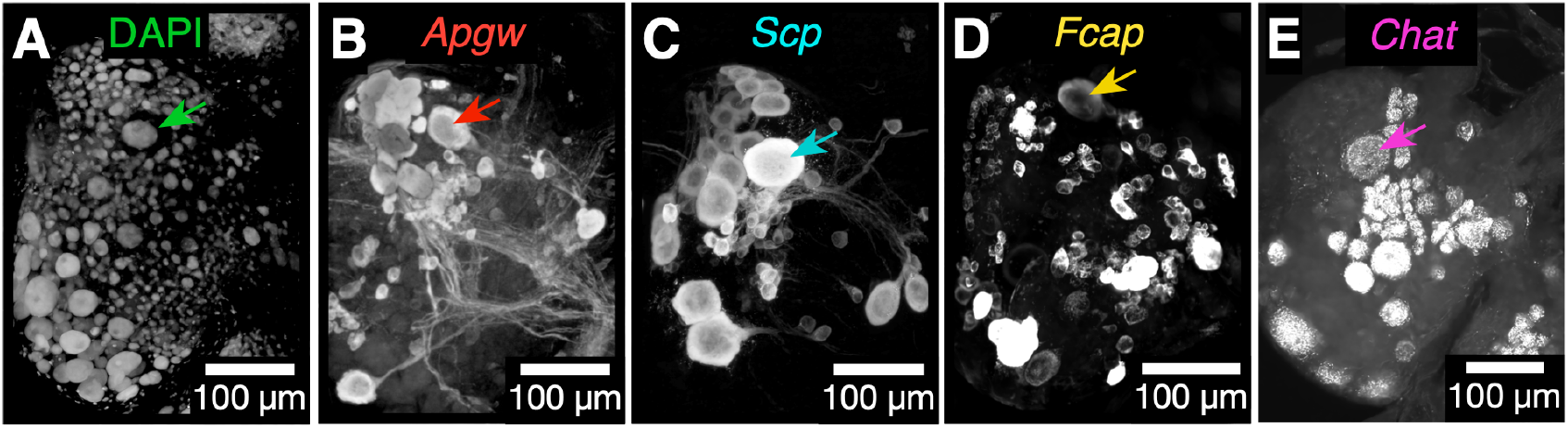
Molecular fingerprint for the giant ventral peptidergic neuron was assembled by analyzing different images of gene expression patterns with Z-projection of a fluorescence confocal image using HCR. Each image shows the right *ceg* and *plg*. Arrows indicate the large identifiable neuron in each. A) Nuclei labeled with DAPI. (B-E) HCR labeling for: *APGWamide* (B), *Scp* (C), *Fcap* (D) and *Chat* (E). All panels are from different samples.

### Unannotated genes are key to differentiating neuron clusters/groups/types

One striking feature of the top markers for all clusters was the abundance of unannotated genes that were differentially expressed (Fig. 14). Unannotated HOGs represented 38% of the total number of HOGs. The high percentage of unannotated HOGs highlights the historical lack of molecular and functional data to characterize these clade-specific genes found in gastropods and other molluscs. Of unannotated HOGs, only 13% (2,448) were shared among *Berghia* and at least one other gastropod, and the remaining 87% (15912) of unannotated HOGs contained only *Berghia* sequences.

**Figure 14.**
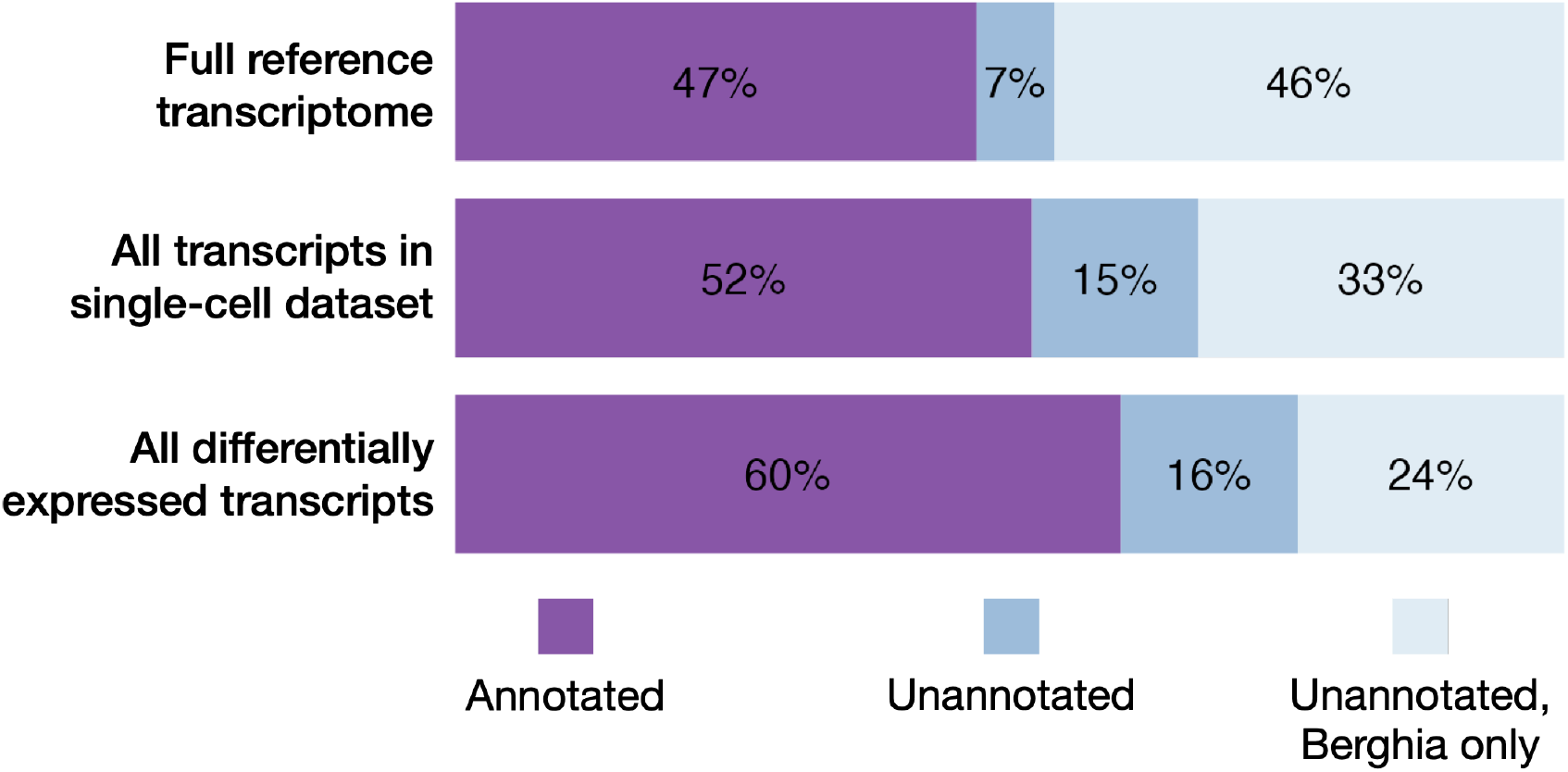
Stacked bar plots of the proportions of Hierarchical Orthogroups (HOGs) for different subsets of the transcriptome that were annotated, unannotated and shared between *Berghia* and at least one other species, or unannotated and *Berghia*-specific. The threshold for differentially expressed genes was a FDR adjusted p-value of less than 0.05.

Out of a total of 24,762 transcripts that ended up in the single cell dataset, 48% (11891) were unannotated, and 52% annotated (12871). Within unannotated transcripts, 31% (3680) were shared between *Berghia* and at least one other species, and 69% (8211) were *Berghia* only. After removing genes with an adjusted p-value above 0.05, there were 3181 differentially expressed genes among all clusters. Forty percent (1279) of differentially expressed genes were unannotated, 39% (500) of which were shared between *Berghia* and at least 1 other species, and 61% (779) were from *Berghia* alone. Unannotated genes are expected for molluscs, where there is much less molecular data and functional characterization of these genes.

We performed HCR for a small selection of unannotated DEGs. As described above, the gene *Bs0019097* was found broadly expressed in putative neuronal clusters in the single cell dataset, and the expression pattern of this gene in most cells that were neurons (see Fig. 4) supports the pattern found in the single cell atlas (see Fig. 3).

*Tph*-expressing serotonergic neurons formed a very small but distinct cluster in the atlas, and the expression patterns for genes in these cells, including *Tph*, *Unc-4*, and *Scf46m3* were validated using HCR (see Fig. 12). Expression of an unannotated gene from the list of DEGs for serotonergic neurons, *Bs0381707*, was restricted exclusively to *Tph*-expressing neurons, as expected for a top cluster marker gene (Fig. 15A,C).

**Figure 15.**
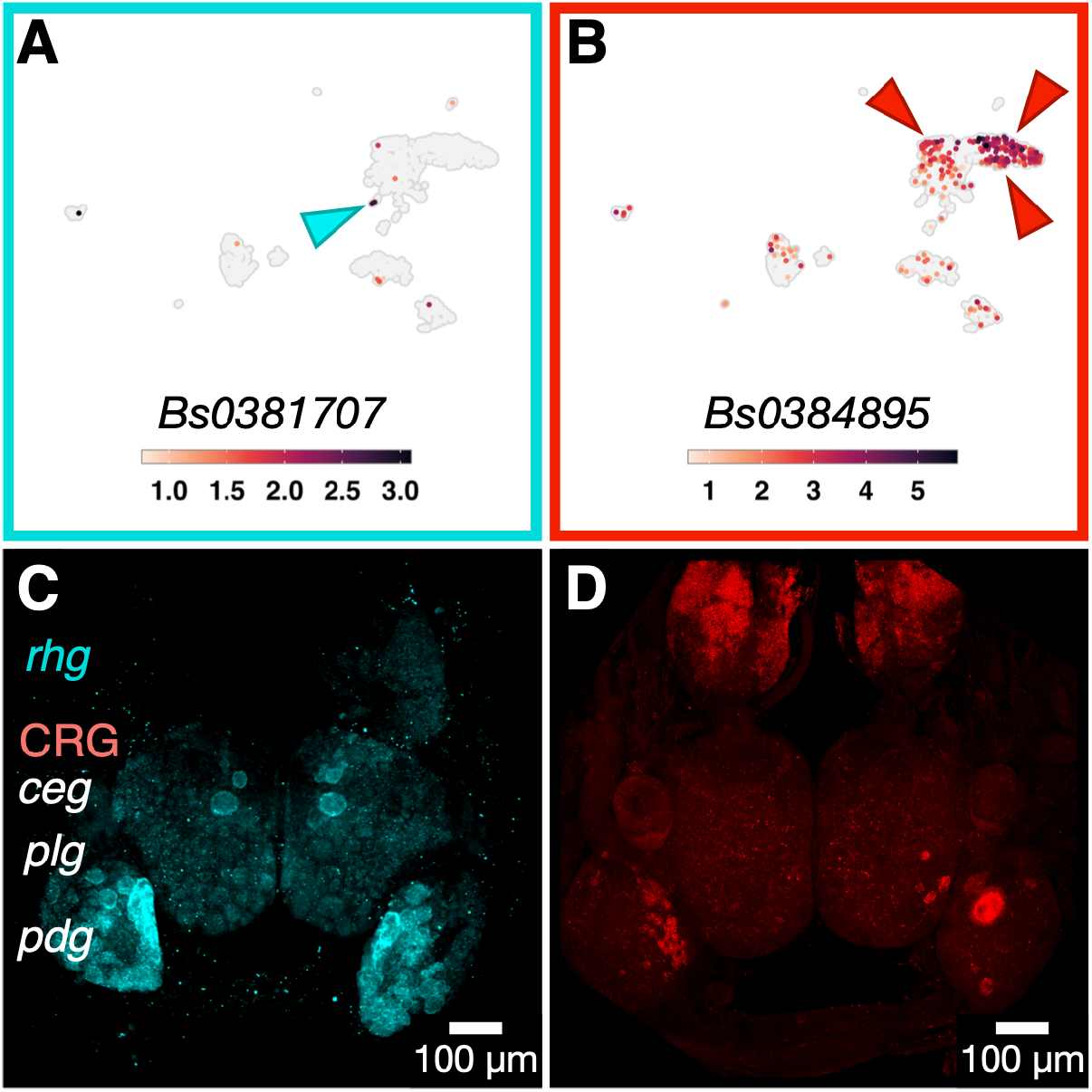
Differentially expressed, unannotated genes are expressed in a cluster-specific manner. A) UMAP plot for unannotated gene *Bs0381707* shows highest expression in the small cluster of *Tph*+ neurons in the single cell dataset. B) UMAP plot for unannotated gene *Bs0384895* shows highest expression in *rhg* clusters in the single cell dataset. C) Z-projection of a fluorescence confocal image using HCR to label mRNA for *Bs0381707* shows its expression was exclusive to the serotonergic, *Tph*+ neurons in the brain. D) Z-projection of a fluorescence confocal image using HCR to label for *Bs0384895* shows widespread, high expression in the *rhg*. However, there were also multiple, small, discrete groupings of *Bs0384895*+ neurons in the *plg* and *pdg*, including one giant neuron in the *pdg*.

Another unannotated gene, *Bs0384895* was differentially expressed in *rhg* neurons (Fig. 15C). Visualization of mRNA for this gene showed high, specific expression in neurons in the *rhg* (Fig. 15D). A small number of neurons outside of the *rhg* also express this gene, including a giant, unilateral, individually identifiable neuron in the right *pdg* (Fig. 15D). These validated examples, as well as the large percentage of unannotated, yet differentially expressed genes in the single cell atlas, support the idea that while unknown, these genes are important for neuronal function and deserve further study.

## Discussion

Individually identifiable neurons are a hallmark of the gastropod CRG, including those of *Berghia stephanieae*. Soma size and location, axonal projections, physiological properties, and gene expression signatures are all measurable phenotypes that can be used to assign individual identities to neurons. Yet, the molecular and developmental mechanisms by which neurons gain their distinct identities in gastropods is unknown. Similarly unknown are the genes that together make up the final mature molecular phenotypes for these neuron types. Only a small percentage of the neuronal population has been identified due in part to the inefficiency of methods previously used to characterize them. It is not currently known whether every neuron, or only a small subset, has an addressable molecular identity. Moreover, the focus on individual neurons has completely ignored whether there is ganglionic organization that could be revealed by molecular signatures of large numbers of neurons. Our study represents an important set of data necessary to begin addressing these questions in molluscan nervous systems.

Using a new study species, *Berghia stephanieae*, we created an annotated reference transcriptome and 1580 single cell transcriptomes from the CRG, and the *rhg*. Standard clustering and differential gene expression analyses generated lists of candidate genes that were explored using multiplexed *in-situ* HCR. We found that there are several broad classes of neurons, some of which are regionally restricted and others of which are scattered throughout the brain. This approach allowed us to map the locations of neurons based on neurotransmitter or neuropeptide phenotype without the concern about cross-reactivity of antibodies yielding false positives. The single-cell resolution of transcriptomic data also revealed candidate DEGs that were not previously known from other transcriptomic analyses of gastropod nervous systems. The results yielded several new insights into genes important for defining neuronal cell type identities and the molecular organization of gastropod nervous systems.

### Glutamatergic and cholinergic neurons are widespread throughout the head ganglia

Neurons expressing markers for small molecule neurotransmitters (e.g., glutamate, acetylcholine, and GABA) were found in all the scRNAseq neuronal clusters. HCR for *Vglut* and *Chat* showed widespread distribution of neuronal somata in all ganglia, and of varying soma sizes. Thus, these two neuronal classes are large umbrella classes containing high numbers of different neuronal cell types. Glutamate and acetylcholine are likely to play major roles in neural circuits in *Berghia* as has been suggested by pharmacological studies in other nudibranchs (***Megalou et al., 2009***; ***Sakurai and Katz, 2017***). In the nudibranch *Hermissenda*, both acetycholine and GABA have been suggested as neurotransmitters in eye photoreceptors (***Heldman et al., 1979***; ***Schultz and Clark, 1997***). Contrasting with these previous findings, *Vglut*, but not *Chat* or *Gad*, was expressed in eye photoreceptors. Both *Vglut* and *Chat* were expressed in neurons in the optic ganglia. *Gad* mRNA was found in a small number of neurons in the scRNAseq atlas data as well as in the brain itself. *Gad* expression was consistent with previous studies of GABA IHC which showed a more restricted distribution across neurons in nudibranchs (***Gunaratne et al., 2014***; ***Gunaratne and Katz, 2016***).

### Serotonergic neurons form a distinct class with multiple neuronal cell types

In contrast to markers for glutamate, acetylcholine, and GABA, markers for serotonergic neurons were highly restricted to a single UMAP cluster and the somata were found in specific locations in the *ceg* and *pdg*. This distribution of somata was known previously from serotonin IHC in other nudibranch species (***Newcomb et al., 2006***). Many of these neurons have been studied for their roles in motor behaviors in other gastropods (***Getting et al., 1980***; ***Jing and Gillette, 1999***; ***Lillvis et al., 2012***; ***Lillvis and Katz, 2013***; ***Yeoman et al., 1994***; ***Zhang et al., 2003***).

Because they formed a discrete cluster in the scRNAseq atlas, we were able to look for differentially expressed genes that were specific to serotonergic neurons. We found that all serotonergic neurons expressed *Unc-4*, but *Unc-4* expression was not restricted to those neurons (Fig. 12). In contrast to our results from *Berghia*, *Unc-4* is primarily associated with cholinergic motor neurons in both *C. elegans* and *Drosophila* (***Lacin et al., 2020***; ***Pflugrad et al., 1997***). This raises at least two potential evolutionary scenarios: *Unc-4* was co-opted from cholinergic neurons to also specify serotonergic neurons or the neurons specified by *Unc-4* switched neurotransmitter phenotype. Because cholinergic neurons in *Berghia* are not also specified by *Unc-4*, the first scenario of co-option may be less likely. Therefore, an ancestral coding of an efferent neuronal type may have undergone a transmitter shift since the common ancestor of protostomes.

### Different patterns for neuropeptide expression

We were able to recover evidence for expression of 40 neuropeptides, 35 of the 65 known molluscan neuropeptide families, which was unexpected given the small total number of single cell transcriptomes in the dataset. High complexity in our small sample of neurons suggests that many neuropeptides may be expressed in many different neuron types rather than being restricted, for the most part, to a single neuronal class. More extensive surveying may find combinatorial expression of multiple neuropeptides in each neuron as seen in the giant ventral neuron, which expressed at least three neuropeptides: *APGWamide*, *Scp* and *Fcap* (Fig. 13). The survey of expression patterns for 8 neuropeptides showed relatively little overlap of sampled neuropeptides in the same neurons (Fig. 9), again consistent with the high complexity of potential co-expression given the large number of expressed neuropeptides.

### Olfaction-associated genes show spatial organization in the rhinophore ganglia

Neurons in two *rhg* clusters expressed *Nitric oxide synthas*e (*Nos*). Nitric oxide (NO) signaling is used by neurons in the olfactory centers of many animals, including molluscs like the snail *Lymnaea stagnalis* (***Gelperin, 1994***). As nudibranch rhinophores are thought to be distance chemoreceptors, expression of *Nos* in the *rhg* is consistent with that function. Outside of the *rhg*, *Nos* was also found in some neurons in the *ceg* and *plg*. This is a broader distribution than would be expected by previous studies using IHC or NADPH diaphorase staining in other nudibranchs (***Hurst et al., 1999***; ***Moroz and Gillette, 1996***; Newcomb and Watson, 2001).

The receptor for NO, *Soluble guanylate cyclase* (*Sgc*) (***Martin et al., 2005***) was differentially expressed in the scRNAseq atlas in a separate cluster of *rhg* neurons than those expressing *Nos*. Multiplexed HCR for *Nos* and *Sgc* in the *rhg* revealed the spatial separation of *Nos* and *Sgc*-expressing populations of neurons. This spatial organization of *Nos*, *Pdf*, and *Sgc*-expressing neurons in the *rhg* hints at complex structural organization of the *rhg*, a ganglion that has received significantly less attention than the CRG. This complexity would not have been apparent without both the ability to detect differential gene expression between populations of neurons using high-throughput methods, and the ability to easily visualize the mRNA within those groups of neurons via HCR.

The expression of other olfaction-associated marker genes in the *rhg* is also consistent with the expected function of the rhinophores as distance chemoreceptors. One *Nos*-expressing *rhg* cluster in the scRNAseq atlas showed differential expression of the neuropeptide *Pigment-dispersing factor* (*Pdf*). Consistent with the separation seen in the scRNAseq atlas data, multiplexed HCR labeling of *Nos* and *Pdf* found populations of *Nos*-expressing *rhg* neurons that were spatially segregated from those that expressed both *Nos* and *Pdf*. In the snail *Helix*, *Pdf* is expressed in multiple parts of the CRG, including the procerebrum, thought to be their olfactory center (***Elekes and Nässel, 1999***). *Enterin* is also differentially expressed in *rhg* clusters in the scRNAseq atlas. In the garden slug *Limax*, *Enterin* is expressed in their olfactory centers, the procerebrum and tentacular ganglia. Ectopic application of *Enterin* modulates local field potential oscillations in these ganglia (***Matsuo et al., 2020***).

### Expression of transcription factors (TFs) also reveals molecular organization at different spatial scales within the head ganglia

There are regional and ganglionic expression differences in transcription factors. Expression of *Six3/6*, a member of the *Six/sine oculis* family of homeobox containing TFs, is primarily restricted to the anterior-most major head ganglia (the *ceg* and *rhg*). This expression pattern is shared across bilaterians, including mammals (***Conte et al., 2005***), indicative of conservation of an ancestral role for *Six3/6* in delineating anterior portions of nervous systems.

The activity of TFs is usually associated with developmental processes in animals, but they also act as “terminal selectors” whose expression maintains mature neuronal phenotypes (***Hobert and Kratsios, 2019***). For example, despite its canonical role in early development, expression of *Delta* in mature neurons in adults has also been described in *Drosophila* and vertebrates (***Cornbrooks et al., 2007***; ***Stump et al., 2002***). In *C. elegans*, *Delta* (*lin-12*) expression also influences locomotor behavior without changing cell type identities, indicating functions beyond development (***Chao et al., 2005***). Both *Six3/6* and *Delta* were DEGs in the glutamatergic *Sgc rhg* neuron cluster of the scRNAseq atlas. Multiplexed HCR showed that they were often co-expressed in *rhg* neurons. Yet *Six3/6* and *Delta* expression was not restricted only to the *rhg*, further supporting their role as a broad neuronal class marker.

Many other transcription factors were found in the scRNAseq data. Those TFs associated with neuroblasts and newly differentiated neurons like *Sox2*, *Sox6*, and *Scratch* were differentially and specifically expressed in one cluster in the scRNA-seq atlas. It is not yet clear whether there are neuroblasts in the brain itself—neurogenesis zones in other gastropods are thought to be in the body wall epithelium, with postmitotic neurons migrating into the CRG from the periphery (***Jacob, 1984***).

### The prevalence of unannotated genes

“Conserved hypothetical proteins” (***Galperin and Koonin, 2004***; ***Rocha et al., 2023***) have emerged as important foci for functional research. The importance of looking at these types of unannotated genes was highlighted by this project; unannotated genes made up approximately 40% of DEGs between clusters, both neuronal and non-neuronal. Although we did not specifically analyze the lineage conservation of unannotated genes in our dataset, we explicitly considered the conservation of *Berghia* sequences with at least one other gastropod as part of the filtering of the reference transcriptome.

The expression of two such markers were visualized using HCR and the expression patterns corresponded specifically to their respective clusters. For example, *Bs0381707* is an unannotated gene shared among nudibranchs, and in *Berghia*, is only expressed in serotonergic neurons (Fig. 15A,C). Another unannotated gene, *Bs0384895*, is highly expressed in *rhg* neurons and in a very restricted set of *pdg* neurons (Fig. 15B,D). While not a cluster-specific marker, unannotated gene *Bs0019097* was widely expressed in putative neuronal clusters and labeled neurons on all ganglia (Fig. 4). The exact nature of *Bs0019097* remains unknown, its distribution and co-expression with genes like *Elav-1* and *7B2* establishes it as a pan-neuronal marker.

The molecular genetics of gastropods has not been well characterized because of the difficulty in genetic manipulations to test gene function in these animals. Some unannotated genes were expected based on the number of “hypothetical proteins” available in molluscan genomes, including a recent chromosome-level genome for *Berghia stephanieae* (***Goodheart et al., 2023***), and recent scRNAseq studies have also described unannotated genes as markers for specific populations of neurons in other molluscs (***Albertin et al., 2015***; ***Songco-Casey et al., 2022***; ***Styfhals et al., 2022***). The great number of unannotated genes shows how much there is to learn about these molecules and their functions in the nervous system.

### Gene expression profiles of individual neurons

Because individual neurons can be identified in gastropods (***Croll, 1987***; ***Katz and Quinlan, 2019***; ***Leonard, 2000***), we expected to find that neurons would have clear molecular signatures. However, the methodology used here was biased against finding rare cells that did not share gene expression profile similarities with many other cells. That said, we were able to show that one neuron, the giant ventral peptidergic neuron, which is visually identifiable based on soma size and position, did have a particular signature. We found that this neuron expressed at least three neuropeptides and *Chat*. This suggests that intersectional, combinatorial and spatial transcriptomic analyses might provide gene expression fingerprints for other uniquely identifiable neurons.

### Functional implications of gene expression

Suites of co-expressed genes that may represent evolutionarily conserved neuronal types were identified in the *Berghia* single cell dataset. For example, the co-expression of *Brn3* and *Vglut*, which was found in *Berghia*, suggests that these cells may be part of mechanosensory circuits, based on the known bilaterian molecular signature for mechanosensory-related neurons. It also suggests that these neurons are potential homologs are of mechanosensory neurons in *Lymnaea* and *Aplysia* (***Brunet Avalos et al., 2019***; ***Nomaksteinsky et al., 2013***). However, we did not find extensive co-expression of *Brn3* and *Sensorin-A* as was reported in S-cells / J,K cluster cells in other gastropods (***Nomaksteinsky et al., 2013***). Instead, a few cells adjacent to the S-cells expressed *Brn3*. There were a small number of neurons that did co-express these genes, but they were located at the lateral edges of the ganglia and clearly not part of the S-cell cluster.

### Multiple glial cell types, including giant glia

Four types of glial cells were previously identified in other gastropods based on immunoreactivity to glial fibrillary acid protein (Santos et al., 2002). We identified three clusters of glial cells in the scRNAseq data. A differentially expressed gene shared among the 3 glial clusters was *Apolipophorin*, which has been identified as a glial marker in recent single cell papers from other molluscs (***Styfhals et al., 2022***). *Apolipophorin* is known to be expressed in *Drosophila* astrocytes (***Yin et al., 2021***), strengthening its use as a glial marker in molluscs. HCR for *Apolipophorin* mRNA revealed a multitude of cells that are likely glia, including a handful of individually identifiable, giant encasing glial cells. Giant glia have been reported previously in leech ganglia (***Deitmer et al., 1999***). In leech, these segmental glia encase all neuronal soma. Here, we found that giant glia encased numerous neuronal soma, but the majority of neurons were associated with other, smaller glia, and not the giant glial cells. We do not yet understand the nature of the relationship between giant glia and the neurons they encase, nor whether there is a typal or functional significance to which neurons are encased.

### Summary

Molluscs, and specifically gastropods, have been important study systems in neuroscience for decades from a cell and circuit perspective. This project shows that new egalitarian molecular tools can be applied to gastropods like *Berghia* to accelerate our understanding of the molecular bases for neuronal identity and function. This will expand the role for gastropods in modern neuroscience. Gastropods, by virtue of their phylogenetic position as molluscs, provide an important counterpoint to other standard laboratory species to help establish general principles of nervous system structure and function.

## Methods and Materials

### *Berghia* husbandry

Adult *Berghia* were originally purchased from online suppliers Salty Underground (Crestwood, MO, USA) and Reef Town (Boynton Beach, FL, USA). They were housed in 5-gallon glass tanks with about 10 individuals. Multiple sea anemones (*Exaptasia diaphana*) were provided to each tank every other day as food. Colonies of *Exaptasia* were raised in 10-gallon tanks with continuously running filters and were fed every other day with freshly hatched *Artemia nauplii*, which were hatched every 2 days by placing 2.5g of freeze dried *Artemia* eggs into an aeration chamber with fresh artificial seawater (Instant Ocean Spectrum Brands, Blacksburg, VA). All animals were kept in the same room, which was held at 26°C to mimic conditions in *Berghia*’s ambient environment in the Florida Keys, USA. The room that housed all the animals was kept on a constant 12:12 light: dark cycle. Larger (1^∼^1.5cm), egg-laying adults were chosen from random tanks for dissection.

### Single neuron RNA sequencing

Brains were dissected and treated in a mixture of 2% w/v pronase (Sigma-Aldrich, St. Louis, MO) and 0.2% w/v Liberase (Sigma-Aldrich, St. Louis, MO) in calcium-, magnesium-free artificial sea-water (CMFSW) to digest the ganglionic sheath. The rhinophore ganglia (*rhg*) were removed with scissors from the CRG at the connective. A total of 18 *rhg* pairs and 20 CRG samples were recovered. *rhg* and CRG samples were separately pooled to create one sample of each. The two samples were processed separately because of a large difference in the average size of the neurons; most of the *rhg* neurons were less than 10*μ*m in diameter, whereas CRG neurons ranged from 10-60*μ*m in diameter.

Each sample was triturated in a 400*μ*L of CMFSW until a single-cell suspension was achieved with minimal cell clumps. CRG samples were mixed for 2 minutes with regular-bore P200 tip at a quick pace, 1 minute with wide-bore P200 tip at a slower pace, and 1 minute with regular-bore P200 tip at the same slow pace. *rhg* samples were mixed for 4 minutes with regular-bore P200 tip at a quick pace. Cell suspensions were filtered through a 400*μ*L layer of 4% bovine serum albumin (BSA) in CMFSW at 4°C. The suspensions were then centrifuged in a swing bucket centrifuge at 4°C for 10 minutes at 100 x g for the CRG sample and for 6 minutes at 400 x g for the *rhg* sample.

After centrifugation, the cell pellets were resuspended in 1.5mL round bottom tubes (Eppendorf DNA LoBind, Eppendorf, Enfield, CT) with 400*μ*L of CMFSW and fixed in 1.6mL of 100% methanol. Precipitants form at this step from the salt in CMFSW. We incubated tubes for 10 minutes at -20°C and centrifuged them in a swing bucket centrifuge for 5 minutes at 4°C and 500 x g. Two mL of 0.5% w/v BSA in 1X phosphate-buffered saline (PBS) was added to dissolve the salt. The suspensions were centrifuged again as above. 400*μ*L of 0.5% w/v BSA in 1X PBS was added first then 1600*μ*L of 100% methanol were added drop by drop to fix the cells. The tubes were stored at -20°C overnight or for several weeks while additional samples were collected.

Single neuron library preparation and high-throughput sequencing cell suspensions were sent to the Bauer Core Facility at Harvard University for library preparation using the 10x Genomics Chromium platform. They also sequenced the single cell libraries on an Illumina NovaSeq.

### Bulk brain transcriptome library preparation and sequencing

*Berghia* brains (CRG plus *rhg*) were dissected and put into lysis buffer from the Smart-Seq4 kit (Takara Bio USA Inc., San Jose, CA) and stored at -20°C. RNA was extracted from the brains and library prep performed following manufacturer’s protocols. We used a magnetic mRNA isolation kit (New England BioLabs, Ipswich, MA) to enrich samples for mRNA before proceeding to library preparation. Prepared libraries yield was quantified using a Qubit dsDNA HS Assay Kit (ThermoFisher Scientific, Waltham, MA) and library quality using the Agilent 2100 Bioanalyzer RNA 6000 Pico assay (Agilent Technologies, Inc., Santa Clara, CA). Some samples could not be quantified because the concentration was too low, but Bioanalyzer traces suggested a quality library, and so these samples were kept in the dataset. Bioanalyzer and bulk transcriptome sequencing was performed by the Genomics Resource Facility at the University of Massachusetts-Amherst using an Illumina Next-Seq 500.

### Brain transcriptome assembly, phylogenetic validation of transcripts and annotation

The raw reads of the bulk brain samples as well as samples from tissues across the body of *Berghia* downloaded from NCBI (SRX10690963, SRX10690964, SRX10690965, SRX10003038, SRX10003039, SRX8599769, SRX8599770, SRX8599771, SRX8599772, SRX8599773, SRX8599774, SRX8599775) were concatenated and put into the Oyster River Protocol (ORP) (***MacManes, 2018***) pipeline for assembly. This pipeline uses multiple assemblers and multiple k-mers. This is necessary because different algorithms assemble true transcripts that are missed by others (***Smith-Unna et al., 2016***). The pipeline uses trimmomatic (***Bolger et al., 2014***) to trim reads to remove adapter sequences and low-quality reads. The cleaned reads are assembled using Trinity (***Grabherr et al., 2011***), Oases (***Schulz et al., 2012***) and SOAPdenovo (***Xie et al., 2014***). Orthofinder2 (***Emms and Kelly, 2018***) has been modified to concatenate sequences across transcriptomes from different assemblers to reduce duplicate sequences. We used this same pipeline to assemble new brain transcriptomes for other nudibranch species. Raw reads for the *Dendronotus* brain transcriptome were uploaded to NCBI (BioProject PRJNA1009839). *Hermissenda* and *Melibe* brain raw reads were downloaded from NCBI (SRX811408; SRX1889794).

We used a phylogenetic approach to further consolidate the *Berghia* transcriptome and clear out spurious transcripts. We assumed that if a homologous transcript could be found in at least 2 other gastropod species, that it is a real transcript. Orthofinder2 was used to create a list of *Berghia* transcripts that met those criteria. Orthofinder2 takes transcriptomes from multiple species and attempts to assign transcripts to hierarchical orthogroups (HOGs). Orthogroups are first created using diamond (***Buchfink et al., 2021***) to search and group transcripts from each species using sequence similarity, MAFFT (***Katoh and Standley, 2013***) to align the sequences per orthogroup and FastTree (***Price et al., 2010***) to build a gene tree for each orthogroup. Orthofinder2 then uses the outputs of this first step to find single copy orthologs and create a species tree. Finally, Orthofinder2 performs a gene-species tree reconciliation for each orthogroups’ gene tree to identify the duplications and losses of orthologs and paralogs that can confuse proper assignment of orthology and to place these events on each gene tree.

The output includes a list of each orthogroup, now put into a unique HOG after the gene-species tree reconciliation, and the transcripts from each species that are putative orthologs. We gave Orthofinder2 the coding sequence transcriptomes from gastropod species with genomes, such as *Aplysia*, *Lottia* and *Pomacea*. We also included our *Berghia* transcriptomes as well as brain transcriptomes from multiple other nudibranch species, including *Melibe leonina*, *Dendronotus iris*, and *Hermissenda opalescens* (formerly identified as a sample from *Hermissenda crassicornis*). Finally, we included sequences for neuropeptides known from other lophotrochozoans that have been verified in the marine worm *Platynereis dumerilli* taken from ***De Oliveira et al. (2019)***. Any HOG that did not have a transcript from *Berghia*, or that had transcripts from fewer than two other species of gastropods were excluded from downstream analyses at this stage.

The *Berghia* transcriptome from ORP was annotated using EnTAP (***Hart et al., 2020***), and Trinotate (***Ghaffari et al., 2014***). These annotations, along with those from *Aplysia*, *Pomacea*, and *Platynereis* were then assigned to HOGs, which could then be mapped onto transcripts from *Berghia*. We found this approach necessary because we found many unannotated transcripts were differentially expressed between neurons using our single cell transcriptome data. It was difficult to know whether these represented true transcripts that do not share any similarity with annotated sequences, or chimeric or artifactual transcripts from assembly errors.

Final HOGs were selected based on the following criteria: 1) they include a sequence from *Berghia* and at least one other species or 2) the *Berghia* sequence was predicted as a complete peptide at least 30 amino acids long. While these permissive criteria may keep some sequences that are spurious or artifacts of assembly, we included them because we know that many neuropeptides are fewer than 100 amino acids. Our expectation is that reads from single cell transcriptomes should not map well to spurious sequences. For similar reasons, we kept sequences even if they could not be annotated using any publicly available database.

### Single cell gene expression clustering analysis and differential gene expression analysis

Raw reads for each sample type were processed for quality and adapters trimmed using HTStream (***Petersen et al., 2015***) with default settings. kallisto (***Bray et al., 2016***) was used to pseudo-map reads onto our reference transcriptome and bustools (***Melsted et al., 2021***) to generate gene counts per cell. After mapping, the outputs of bustools were imported into R using the bustools kbk command. Kneeplots were used to exclude empty droplets and those containing ambient RNA. A standard Seurat (***Hao et al., 2021***) normalization and analysis pipeline was used on the single cell transcriptomes. While exploring a preliminary analysis, we discovered that some cells expressing *Tryptophan hydroxylase* were labeled as *rhg*, despite evidence using serotonin IHC that there were no neurons in the *rhg* that were serontonergic (***Whitesel, 2021***). These cells also shared the exact barcode with another cell from the CRG sample within the cluster, which is highly statistically unlikely. We discovered that these “*rhg*” barcoded cells were likely an artefact of multiplexing, which is common to single cell RNA sequencing datasets but not often considered (***Griffiths et al., 2018***). To resolve this issue within the entire dataset and recover cells rather than remove them entirely, similar principles as (***Griffiths et al., 2018***) were used to determine the most likely sample of origin. When a barcode was shared, the number of UMIs (Unique Molecular Identifiers) were counted. There was a clear split for most cells of the proportion of UMIs present from one sample or the other. A barcode was called for either the CRG or *rhg* sample if it contained at least 70% of the total UMIs with the shared barcode. The other cells was removed from the dataset before further processing with Seurat as follows. Briefly, each sample type was imported separately, and genes that were not expressed in at least 3 cells, and cells expressing fewer than 200 genes were excluded. The data were merged while keeping track of the origins of data for each cell and checked for batch effects looking at the number of UMIs and genes in each sample. There was no evidence for batch effects, so all downstream analyses were performed on both samples combined. After removing cells that failed to meet the above criteria, counts for each gene were normalized by the total expression in each cell, scaled by multiplying by 10,000, and finally log-transformed. The FindVariableFeatures command was used to identify the 2000 most highly variable genes.

After running a principal component (PC) analysis, the first 50 PCs were included in dimensionality reductions using tSNE (t-distributed Stochastic Neighborhood Embedding, resolution 3) and UMAP (Uniform Manifold Approximation and Projection) methods as implemented in Seurat. To select the resolution used in tSNE, Clustree (***Zappia and Oshlack, 2018***) was used to look at the effect of changing tSNE resolution on the stability of clusters. The highest resolution value used showed only low instability of cells in clusters compared to the next smallest value. We looked for differentially expressed genes between clusters using the FindMarker command in Seurat, using MAST (***Dal Molin et al., 2017***; ***Finak et al., 2015***). Clusters where differentially expressed genes were mostly the same were collapsed together using the Clustree results. There were 14 clusters in the final analysis. We used the FeaturePlot and DoHeatmap commands in Seurat to visualize gene expression. We also used the R package *nebulosa* (***Alquicira-Hernandez and Powell, 2021***) to estimate the kernel density of expression for genes of interest. SCpubr was used to produce UMAP plots, dot plots and heatmaps (***Blanco-Carmona, 2022***). Figures were composed using Graphic (Autodesk, San Francisco, CA, USA). All code and files used for these analyses are available on Github.

### Creating DNA probes for in-situ Hybridization Chain Reaction (HCR)

BLASTP (***Altschul et al., 1990***; ***Camacho et al., 2009***) was used with queries from *Aplysia* and *Lymnaea* to look for candidate genes in the reference transcriptome. ExPAsy Translate (***Gasteiger et al., 2003***) was used to translate the nucleotide sequence into amino acids for possible reading frames, to determine whether the sequence was the sense or anti-sense strand. If the sequence was anti-sense, the reverse complement of the sequence was identified using the online Sequence Manipulation Suite’s reverse-complement tool (***Stothard, 2000***). The longest predicted peptide was selected and used BLAST on NCBI databases to verify that the best hits for the predicted peptide matched the query sequence where possible. The best BLAST hits needed to be from molluscs and have the same annotation where annotations were available. Sequences for the probe sets for each gene were made using HCRProbeMaker V3 (***Kuehn et al., 2021***) with the sense strand, based on the requirements necessary for HCR found in (***Choi et al., 2018***). This script found a user-selected number of probe pairs that would bind to the target mRNA sequence with 2 base pairs in between and added one of 5 initiator sequences to each partner that are necessary for the initiation of the fluorescent hybridization chain reaction. Probe set oligos from this script were synthesized by IDT (Integrated DNA Technologies, Coralville, IA, USA) and the lyophilized probes were rehydrated with 50 *μ*l TE buffer to end up with a 50 picomolar stock concentration. Probe sets for *Choline acetyltransferase* (54 probe pairs) and *APGWamide* (20 probe pairs), as well as the fluorescent hairpins were purchased from Molecular Instruments (Los Angeles, CA, USA).

### Labeling transcripts using HCR

Hybridization chain reaction reagents were made using the recipes from (***Choi et al., 2018***), substituting urea one-for-one instead of formamide. This substitution has been tested for traditional colormetric *in-situ* hybridizations and shown to be equally effective (***Sinigaglia et al., 2018***). Both the formamide and urea formulations were compared using HCR, without any appreciable difference in results. Urea-based hybridization solutions were used for all HCR to minimize use of hazardous reagents. Samples were fixed for at least 2 hours (at room temperature) to overnight (at 4° C) in 4% paraformaldehyde (Electron Microscopy Sciences, Hatfield, PA, USA) in artificial sea water. After fixation, samples were washed briefly in phosphate-buffered saline (PBS, ThermoFisher Scientific, Agawam, MA, USA) and then dehydrated through a series from 100% PBS to 100% methanol. Samples in methanol were stored at least overnight at -20° C.

On the first day of HCR processing, samples were rehydrated in series into 100% 5X sodium citrate saline (SSC, diluted from 20X stock solution, (Thermofisher Scientific, Agawam, MA, USA) and 0.1% Tween-20 added (SSCT). They sat in 5X SSCT at room temperature for at least 15 minutes before being moved to a tube containing 200*μ*L of hybridization buffer in a heat block set to 37° C to equilibrate for at least 30 minutes. During this time, probe sets for each gene were thawed at room temperature, and a small volume was mixed into 100*μ*L of fresh hybridization buffer. The concentration of probe needed for successful labeling varied between probe sets, from 0.1 picomolar to 1 picomolar. After equilibration in hybridization buffer, as much of the used buffer was removed from the tube as possible while keeping the sample submerged. The probe-hybridization buffer mixture was then added to the tube, and the sample incubated overnight at 37° C.

The following day, the samples were washed in hybridization wash twice for 5 minutes, and twice for 30 minutes at 37° C. During the final 30-minute wash, the appropriate Alexa-dye-tagged hairpins for each initiator were thawed at room temperature in the dark. Haiprins with Alexa 488, 546, 594 or 647 dyes were used to multiplex labeling of 2-3 genes per sample. 2*μ*L of each hairpin pair per initiator were used in a final volume of 100*μ*L amplification buffer. Each hairpin solution was aliquoted into a 0.2mL PCR strip tube. Hairpins were then “snapped” to linearize them by heating at 95° C for 90 seconds followed by 30 minutes of cooling at room temperature in the dark. All hairpins were then added to the appropriate volume of amplification buffer to a final volume of 100*μ*l. The hybridization chain reaction occurred in the amplification buffer for 1 or 2 overnights at room temperature in the dark. The length of amplification time was determined empirically, but in general, highly expressed genes needed less amplification time, and the lowest expressed genes needed the longest time.

After amplification, the samples were washed in 5x SSCT once. The nuclei were labeled using 1*μ*L 300uM DAPI : 1000*μ*L 5x SSCT, rotating in the dark for 1 hour. Brains were mounted onto long coverglass using 2 small strips of double-sided tape (Scotch Brand, 3M, Saint Paul, MN) as spacers and to attach the small coverglass. Vectashield Vibrance (Vector Laboratories, Newark, CA), DeepClear (***Pende et al., 2020***), or a fructose-glycerol solution (***Elagoz et al., 2022***) was used to clear and mount the samples for confocal imaging. Each probeset was tested at least twice to ensure consistent labeling.

### Confocal imaging of multiplexed HCR samples and image processing

Fluorescent confocal images were taken at 20x using the Nikon A1R-25 or A1R-18 at the Light Microscopy Facility at the University of Massachusetts-Amherst. Images were processed in ImageJ to rotate the image as needed, normalize intensity, adjust brightness and contrast, and create scale bars.

## Acknowledgments

We would like to thank Drs. Duygu Özpolat and Ryan Null for creating HCR probeset development tools. Drs. Deidre Lyons and Jessica Goodheart provided valuable input and annotations for the reference transcriptome. Implementation of MHD-clearing, lightsheet imaging was done with help from Drs. Joseph Bergan and Joseph Dwyer. We thank Dr. Adriano Senatore for use of an unpublished *Dendronotus* transcriptome. Fran De Mora Ocana and Ryan Allen Wight produced replicate HCR samples for several gene sets. Single cell library preparation and sequencing was done through The Bauer Core Facility at Harvard University. Bulk sequencing of *Berghia* brain samples was performed by the Genomics Resource Laboratory, University of Massachusetts Amherst. Confocal imaging was performed at the Light Microscopy Facility at the Institute for Applied Life Sciences, University of Massachusetts Amherst. This project was funded by a NSF Postdoctoral Research Fellowship in Biology PRFB 1812017 to MDR, and NIH grants U01-NS108637 and U01NS123972 to PSK.

---

**Figure 6–Figure supplement 1.**
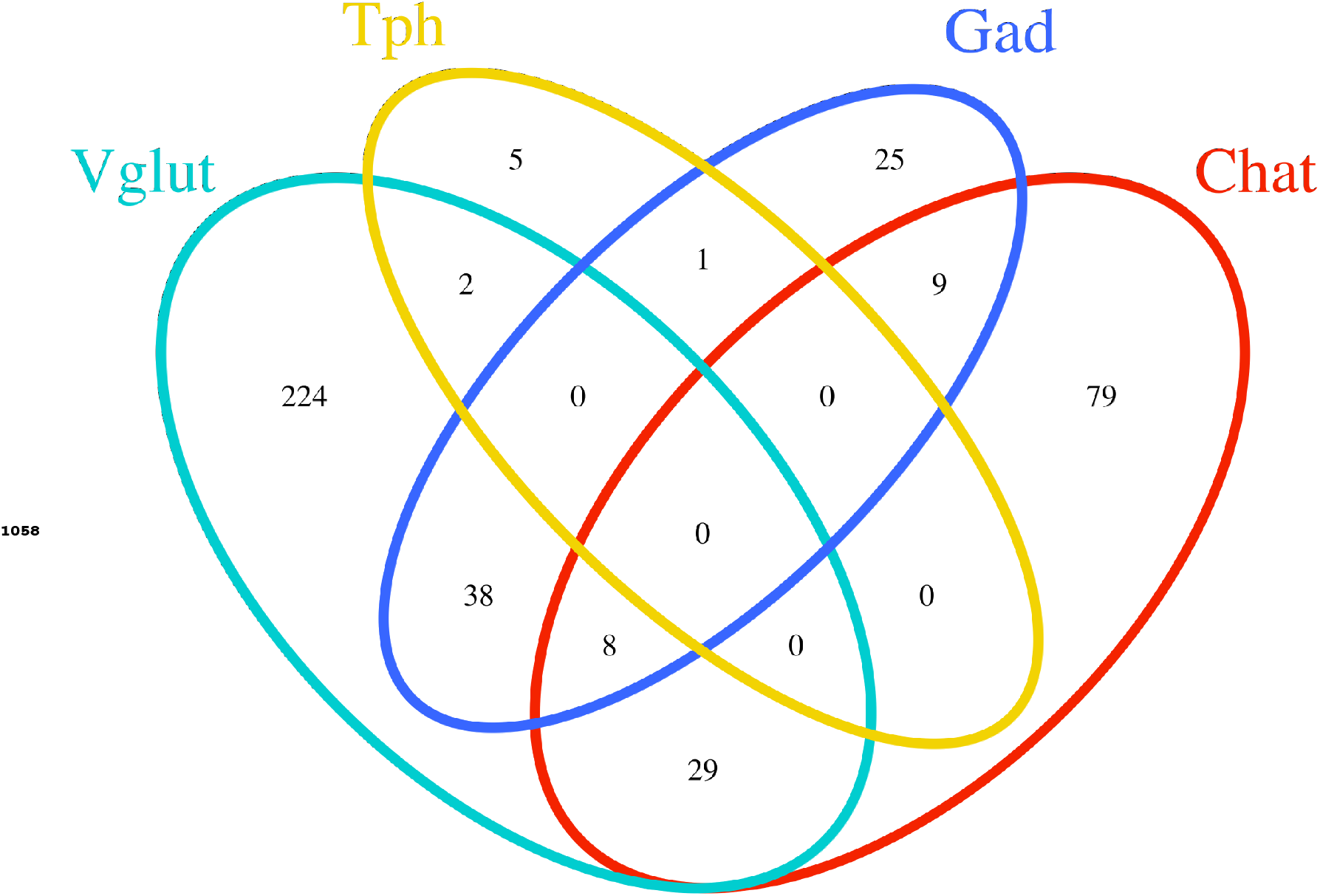
Venn diagram showing the numbers of neurons expressing *Vglut*, *Tph*, *Gad*, and *Chat*, singularly or in combinations. Most neurons express only 1 gene associated with a specific neurotransmitter. *Vglut* shares expression with the largest number of either *Gad*+ or *Chat*+ cells.

