## Supplementary figures and images for "Cellular-resolution gene expression mapping reveals organization in the head ganglia of the gastropod, *Berghia stephanieae*"

### Supplemental Figure 6

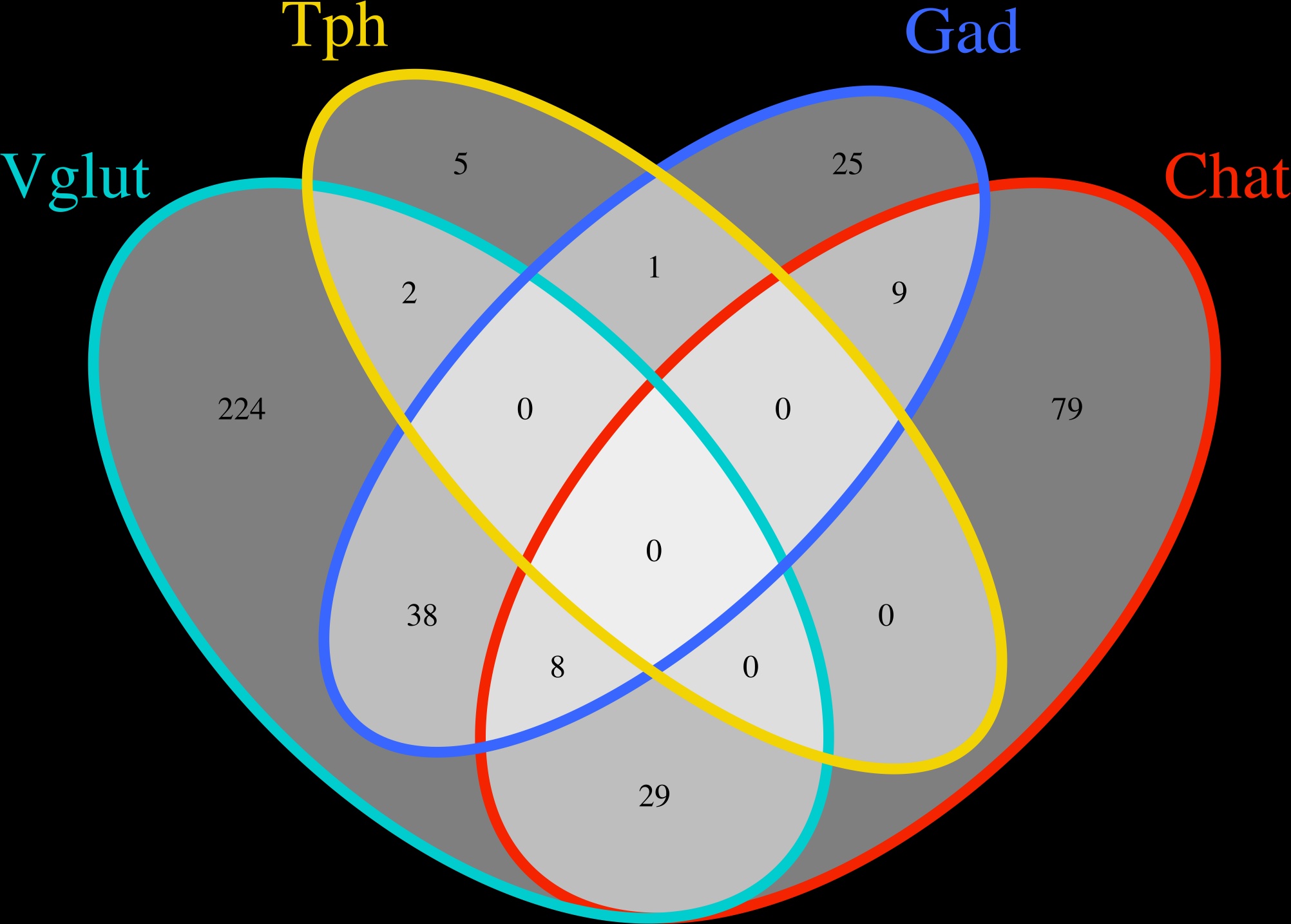
